# The Wg and Dpp morphogens regulate gene expression by modulating the frequency of transcriptional bursts

**DOI:** 10.1101/2020.01.24.918623

**Authors:** Rachael Bakker, Madhav Mani, Richard W. Carthew

## Abstract

Morphogen signaling contributes to the proper spatiotemporal expression of genes during development. One method of regulation of signaling-responsive genes is at the level of transcription. Single-cell quantitative studies of transcription have revealed that transcription occurs intermittently, in bursts. Although the effects of many gene regulatory mechanisms on transcriptional bursting have been studied, it remains unclear how morphogen gradients affect this dynamic property of downstream genes. Here we have adapted smFISH for use in the *Drosophila* wing imaginal disc in order to measure nascent and mature mRNA of genes downstream of the Wg and Dpp morphogen gradients. We compared our experimental results with predictions from stochastic models of transcription. Our results indicate that the transcription levels of these genes appear to share a common method of control via burst frequency modulation. Our data helps further elucidate the link between developmental gene regulatory mechanisms and transcriptional bursting.

## INTRODUCTION

Paracrine signaling is a highly conserved means for cells within a tissue to communicate with one another to regulate diverse activities including proliferation, differentiation, apoptosis, and movement. Many of these activities are mediated by changes in gene transcription that are brought about by reception of the signals. Paracrine factors acting as morphogens are a particularly important class of gene regulators. Morphogens form spatially-extended gradients from the source of their synthesis, and elicit different transcription outputs from target genes, depending on local concentration of the morphogen (Tabata & Takei, 2004).

Many paracrine signals regulate gene transcription via control of the availability or activity of sequence-specific transcription factors. Some transcription factors regulate assembly of the preinitiation complex (PIC) composed of Pol II and general factors at the transcription start site (Esnault et al., 2008). Other factors recruit coregulators that modify nucleosomes or remodel the chromatin architecture of the gene (Bannister & Kouzarides, 2011). However, transcription is a dynamic process, and thus, molecular models of regulation via PIC assembly or chromatin structure, do not adequately capture what kinetic steps in transcription initiation are being regulated. Recently developed methods have uncovered greater complexity in the transcription initiation process than previously imagined. Genes that are constitutively expressed rarely show uniform and continuous mRNA synthesis. Rather, mRNA synthesis occurs in bursts that are interrupted by periods of dormant output. This phenomenon is known as transcriptional bursting (Chen et al., 2019; Chubb, Trcek, Shenoy, & Singer, 2006; Dey, Foley, Limsirichai, Schaffer, & Arkin, 2015; Arjun Raj, Peskin, Tranchina, Vargas, & Tyagi, 2006; Suter et al., 2011).

Various studies have explored how mechanisms of gene regulation affect the size and frequency of transcriptional bursts, and thereby affect transcription output. The availability of transcription factors has been shown to affect burst frequency (Ezer, Moignard, Göttgens, & Adryan, 2016; Larson et al., 2013; Senecal et al., 2014). For example, the *Drosophila* transcription factors Bicoid and Dorsal have been studied in great detail with respect to their effects on transcription burst frequency in the embryo (Garcia, Tikhonov, Lin, & Gregor, 2013; He, Ren, Wang, & Ma, 2012; Holloway & Spirov, 2017; Shawn C. Little, Tikhonov, & Gregor, 2013; Xu, Sepúlveda, Figard, Sokac, & Golding, 2015). Enhancer strength and enhancer-promoter contact correlate with burst frequency of genes (Bartman, Hsu, Hsiung, Raj, & Blobel, 2016; Bothma et al., 2014; Chen et al., 2019; Fukaya, Lim, & Levine, 2016; Larsson et al., 2019). These studies altogether suggest that bursting frequency is potentiated by enhancer-promoter contact and is mediated by transcription factors binding to DNA.

In this study, we have explored how the Wnt protein Wingless (Wg) and BMP protein Decapentaplegic (Dpp) regulate transcription dynamics of genes in the *Drosophila* wing imaginal disc. The Wnt and BMP families of proteins are two highly conserved paracrine factors that can act as morphogens. In canonical Wnt signaling, the binding of extracellular Wnt protein to its transmembrane receptor Frizzled causes β-catenin to be stabilized and free to enter the nucleus, where it relieves repression of Wnt-responsive genes by binding to the sequence-specific transcription factor TCF (Clevers & Nusse, 2012); Swarup & Verheyen, 2012). In canonical BMP signaling, ligand binding to receptor triggers phosphorylation of SMAD proteins, which translocate to the nucleus along with co-SMADs, bind to responsive genes, and activate their transcription (Hamaratoglu, Affolter, & Pyrowolakis, 2014; Shi & Massagué, 2003).

To explore the effects of Dpp and Wg signaling on transcription dynamics, we have adapted single molecule fluorescent in situ hybridization (smFISH) for use in imaginal disc tissues. We use smFISH to quantify nascent and mature mRNAs for several genes expressed in highly diverse patterns within the wing disc. All of the genes are regulated by modulation of transcription burst frequency by Dpp and Wg even though their pression patterns are highly different from one another.

## RESULTS

In this study, we have explored how the Wg and Dpp morphogens regulate transcription dynamics in the wing disc. Each morphogen is synthesized in a narrow stripe of cells within the disc. Wg is produced in cells at the boundary between Dorsal and Ventral (DV) compartments of the wing pouch, while Dpp is produced in cells at the boundary between Anterior and Posterior (AP) compartments (Figure 1A). These factors diffuse from their sources forming concentration gradients across the disc, and regulate gene expression in a concentration-dependent manner.

**Figure 1.**
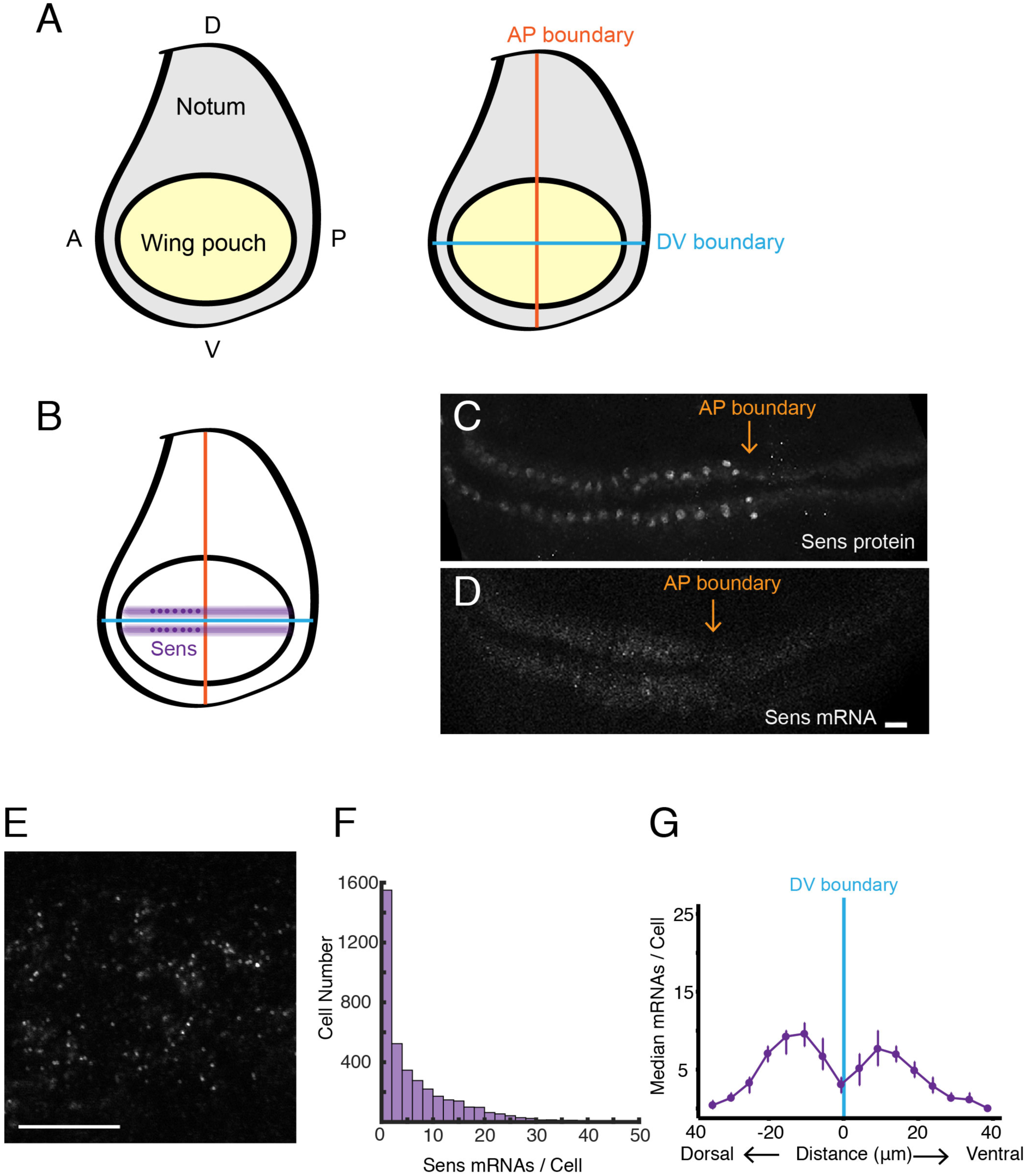
smFISH analysis of sfGFP-sens mRNA levels in wing imaginal discs. **(A)** Schematic of a wing disc outlining different regional domains, and the positions of boundaries between Dorsal - Ventral (V) and Anterior (A) - Posterior (P) compartments of the disc. Each wing disc is composed of roughly 50,000 cells organized in a pseudostratified epithelium. **(B)** Schematized expression pattern for Sens inside the wing pouch centered around the DV boundary. Sens is also expressed in clusters of cells in the notum, which are not shown. **(C-E)** Confocal sections of wing discs expressing sfGFP-Sens. **(C)** sfGFP-Sens protein fluorescence. **(D)** sfGFP-Sens mRNAs as visualized by smFISH using sfGFP probes. Scale bar = 10 μm. **(E)** Higher magnification of sfGFP-Sens mRNAs as visualized by smFISH using sfGFP probes. Scale bar = 10 μm. **(F)** Distribution of wing disc cells as a function of the number of Sens mRNA molecules per cell. **(G)** Sens mRNA number as a function of cell distance from the DV boundary. The shortest path length from each cell centroid to the DV boundary was calculated. Cells were then binned according to this path length and whether they were dorsal or ventral cells. The median mRNA number/cell for each bin is plotted. Error bars represent bootstrapped 95% confidence intervals. A bimodal distribution captures the expression pattern of Sens.

### smFISH detection of mRNA molecules in the wing disc

In order to assay gene expression in the wing imaginal disc, we quantified mRNA numbers using single-molecule fluorescent in situ hybridization (smFISH). With smFISH, a tandem array of fluorescently-labeled oligonucleotides complementary to a given mRNA are hybridized to fixed and permeabilized tissue. When a sufficient number of oligo probes anneal to one mRNA molecule, the aggregate fluorescence can be detected by standard confocal microscopy (A. Raj, van den Bogaard, Rifkin, van Oudenaarden, & Tyagi, 2008). This method has been developed and applied to many systems, including cell culture, *C. elegans*, and the *Drosophila* embryo (Ji & van Oudenaarden, 2012; S. C. Little & Gregor, 2018; Youk, Raj, & van Oudenaarden, 2010). However, it has never been used to probe *Drosophila* imaginal discs, prompting us to develop and refine a robust smFISH method for imaginal discs (see Methods for details).

We first probed for expression of the *senseless* (*sens*) gene in the wing disc. *Sens* is required for cells to adopt a sensory organ fate, and the gene is expressed in two stripes of cells adjacent to and on either side of the DV boundary in the wing pouch (Figure 1B,C). *Sens* expression in the wing pouch is induced by Wg, which is expressed by cells located at the DV boundary (Nolo, Abbott, & Bellen, 2000; Zecca, Basler, & Struhl, 1996). We probed for *sens* mRNAs expressed from a transgenic version of the *sens* gene. We did so for a number of reasons. First, the genomic transgene rescues the endogenous gene based on function and expression (Cassidy et al., 2013). Second, the transgene is tagged such that the amino-terminal coding sequence corresponds to super-fold GFP (sfGFP). By using oligo probes directed against sfGFP, we could easily determine the specificity of detection.

Samples were probed and imaged, revealing the expected pattern of fluorescence localized to two stripes adjacent to the DV midline in the wing pouch (Figure 1D). The fluorescence signal was specific for *sfGFP-sens* since wing discs from larvae not carrying the transgene gave no fluorescence pattern (Figure 1 - figure supplement 1A,B). The fluorescence signal was sufficiently bright that diffraction-limited spots were readily detected in optical sections when imaged under higher magnification (Figure 1E). Each spot was detected in two-to-three contiguous sections, which corresponded to ∼ 230 nm. A custom Matlab script was used to unambiguously identify each of these 3D fluorescent objects (Figure 1 - figure supplement 1C). The frequency distribution of fluorescence intensity for these 3D objects showed a unimodal distribution (Figure 1 - figure supplement 1D). This suggested that the objects had a homogeneous composition, as would be expected for single mRNA molecules. To demonstrate that the fluorescence spots corresponded to mRNA molecules, we incubated wing discs in media containing actinomycin-D, an inhibitor of RNA synthesis. The number of fluorescence spots was greatly diminished (Figure 1 - figure supplement 1E).

To determine the false discovery rate for smFISH detection, we applied two approaches. First, we compared the number of spots in discs expressing the *sfGFP-sens* gene versus discs lacking the gene. From this, we calculated that 0.5 % of identified spots were false-positive (Figure 1 - figure supplement 2A). Second, we co-hybridized *sfGFP-sens* wing discs with two sets of non-overlapping probes - one set recognized *sfGFP* and the other set recognized *sens* sequences. Each probe-set was labeled with a different fluor. If a spot identified using the *sfGFP* probe set was not also identified by the *sens* probe-set, we classified that spot as a false-negative. The analysis indicated that a maximum of 6% of mRNAs (232 out of 3,842 spots scored) were not identified by both probe-sets (Figure 1 - figure supplement 2B). This rate of false-negative identification is comparable to smFISH methods in other systems (A. Raj et al., 2008).

We next wanted to partition identified mRNAs into the cells from which they were expressed. Since the smFISH method denatured the epitopes of all tested antibodies and it also denatured sfGFP, we were unable to segment cells using standard approaches. Instead, we used the fluorescent dye DAPI to visualize cell nuclei in the imaged samples, and we segmented the nuclei into 3D objects (Figure 1 - figure supplement 2C-E). Cell nuclei are located throughout the apical-basal axis of the wing disc because it is a pseudostratified epithelium (Aldaz & Escudero, 2010). Therefore, we approximated the cell boundaries by performing a 3D Voronoi tessellation using the nuclear centroids (Figure 1 - figure supplement 2F). RNAs were then partitioned into the Voronoi cells (Figure 1 - figure supplement 2G).

The abundance of *sens* mRNAs within the DV stripes varied from one to fifty molecules per cell (Figure 1F). This was because the Wg morphogen induces a graded expression pattern of Sens protein across the width of each stripe (Nolo et al., 2000; Zecca et al., 1996). Therefore, we binned cells according to their distance from the DV boundary, and we observed peaks in mRNA number per cell as a function of distance from the boundary (Figure 1G).

We also used the *sfGFP-sens* gene to determine whether the smFISH method could detect mRNAs in other imaginal discs. In the eye disc, *sens* is expressed in a stripe of cells located within the morphogenetic furrow, and indeed we were able to detect smFISH signals in furrow cells of the eye disc (Figure 1 - figure supplement 3). Thus, our method is broadly applicable to imaginal discs.

### smFISH detection of gene expression regulated by Dpp

We extended the analysis to genes downstream of the BMP family protein Dpp. Dpp is expressed in a stripe of cells located at the AP boundary of the wing disc, orthogonal to the Wg stripe (Figure 2A). Dpp protein is transported bidirectionally to form gradients across the disc, and several genes are regulated by Dpp in a concentration-dependent manner. S*palt-major* (*salm)*, *optomoter-blind* (*omb*), *daughters-against-dpp* (*dad),* and *brinker* (*brk*) are expressed in symmetric domains within the anterior and posterior compartments of the wing pouch (Figure 2A,B). *Salm* is symmetrically expressed in a domain somewhat broader than the Dpp stripe, whereas *omb* and *dad* are expressed more broadly, and *brk* is expressed only near the wing pouch border (Celis, Barrio, & Kafatos, 1996; Grimm & Pflugfelder, 1996; Tabata & Takei, 2004). When smFISH was used to detect mRNAs of these genes, it qualitatively recapitulated their known expression patterns (Figure 2C-F). We quantified the number of mRNAs per cell and attempted to map the distribution to cell position within the wing pouch. Since the only landmark we could reliably use was the border between the wing pouch and the rest of the disc, we measured cell position as a function of distance from the border(Figure 2G). When we did so, the distributions in mRNA number per cell gave profiles that were consistent with previous qualitative descriptions of their expression patterns (Figure 2H).

**Figure 2.**
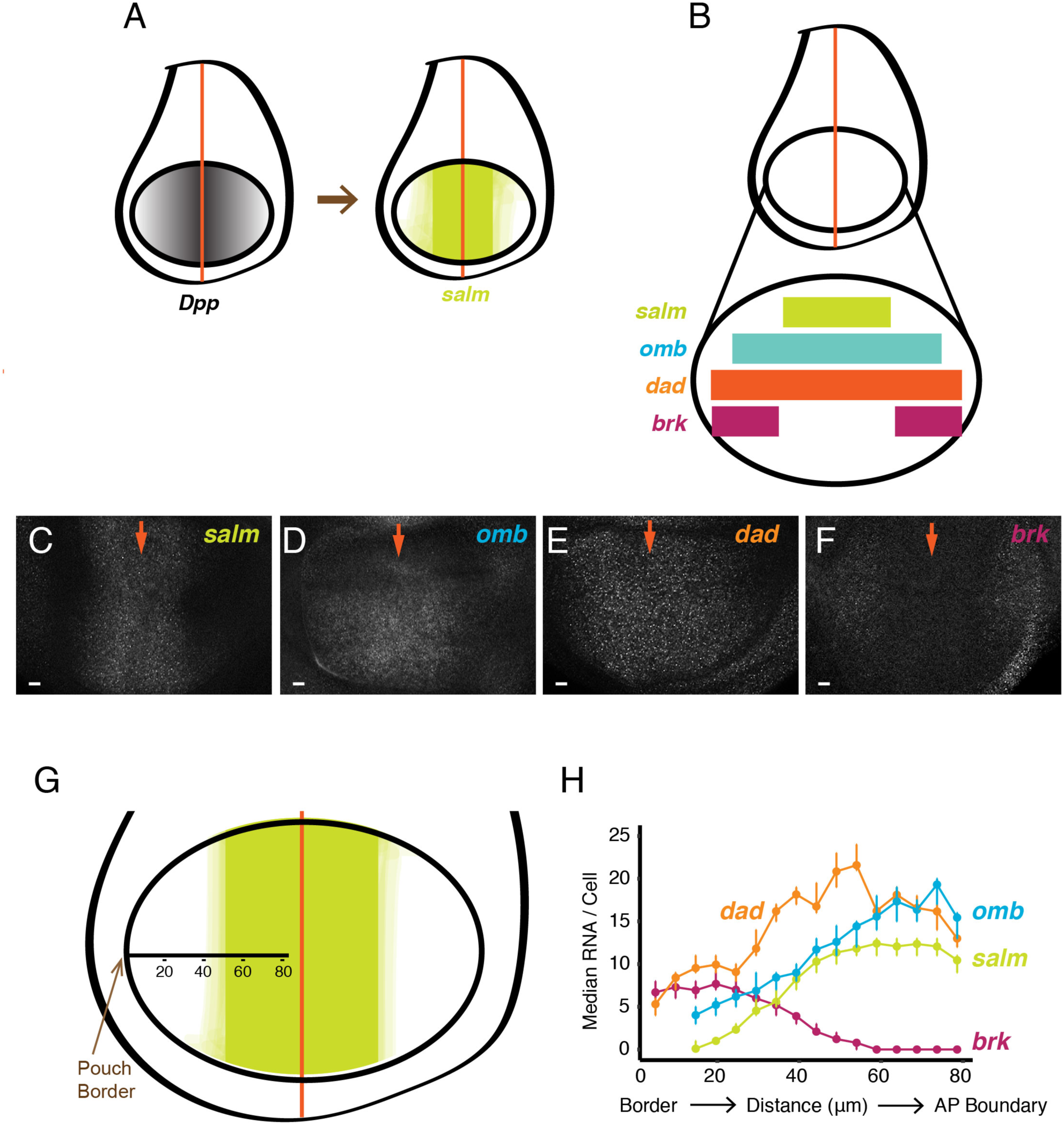
smFISH analysis of mRNA levels from Dpp-responsive genes. **(A)** Schematic of wing discs highlighting the graded distribution of Dpp protein in the wing pouch, centered around the AP boundary, and the expression domain for *salm*, one of the targets of Dpp regulation. Not shown is Dpp localization in the notum domain of the disc. **(B)** Expression domains of four target genes of Dpp signaling. **(C-F)** Confocal sections of wing pouches probed for mRNAs synthesized from the *salm* **(C)**, *omb* **(D)**, *dad* **(E)**, and *brk* **(F)** genes. Orange arrows mark the position of the AP boundary in each image. **(G, H)** mRNA number as a function of cell distance from the anterior-most border of the wing pouch. **(G)** An axis tangential to the AP boundary is used to map cell position. Numbers refer to distance in μm from the wing pouch border. **(H)** Cells were binned according to position along the axis. The median mRNA number/cell for each bin is plotted. Error bars represent bootstrapped 95% confidence intervals.

### smFISH detects sites of nascent transcription

A further benefit to smFISH is that it can detect and quantify RNA as it is being transcribed from a gene. We sought to identify and characterize these sites of nascent transcription in the wing disc. Quantification of pixel intensity of all fluorescent spots revealed two discrete populations: a large population of dim spots of uniform intensity, and a smaller population of brighter spots with more variable intensity (Figure 3A,B). The former population corresponded to those described earlier, and they were primarily located in the cytoplasm - these are the mature mRNAs. The latter population was primarily located inside nuclei, and thus we hypothesized that these were sites of nascent transcription. To confirm that these bright spots corresponded to transcription sites, we used probes complementary to an intron in the *omb* gene. These probes only detected the brighter population of spots localized to nuclei (Figure 3C). Since introns are not spliced out until after transcription, this result supports the conclusion that the brighter nuclear spots are sites of nascent transcription.

**Figure 3.**
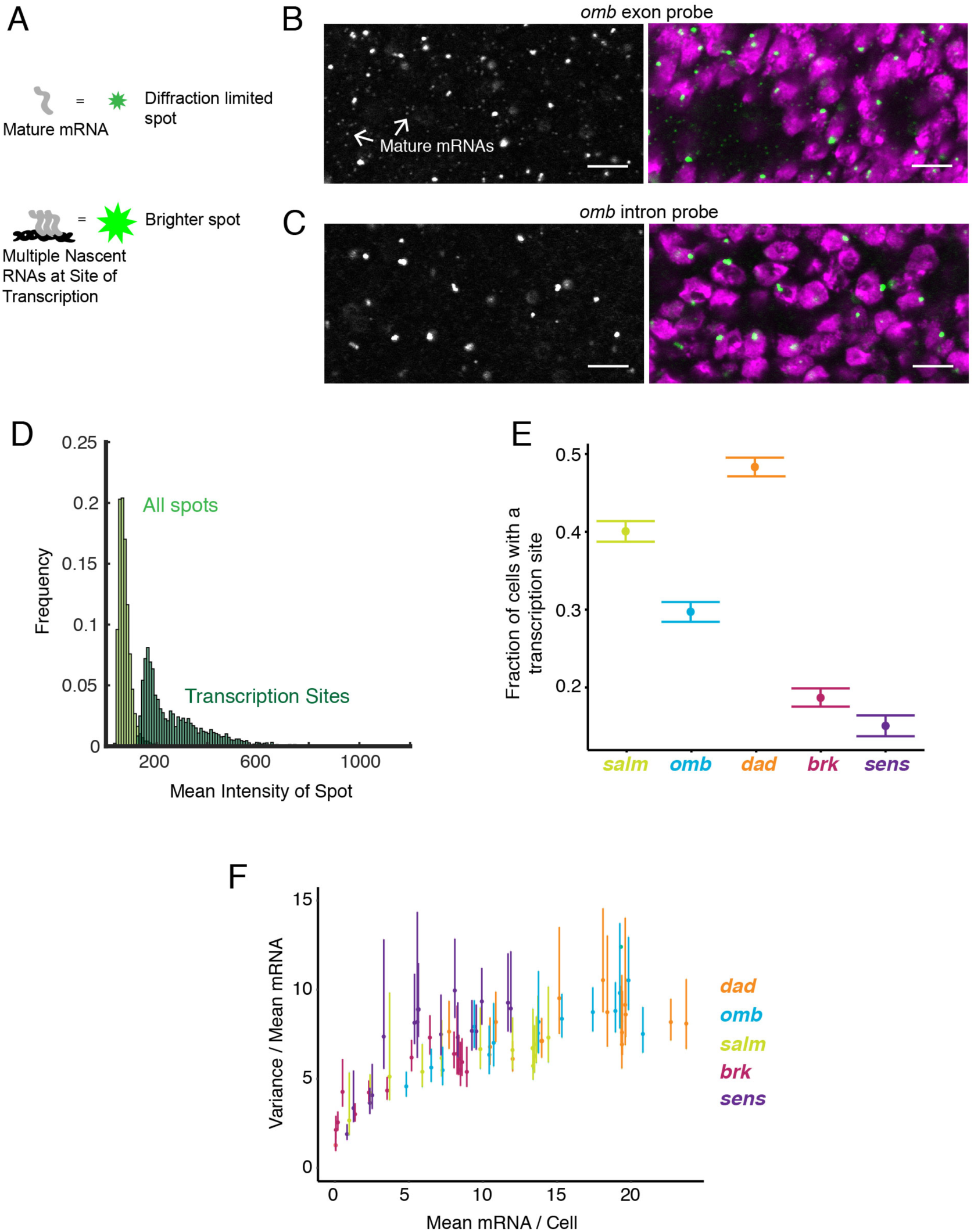
Sites of nascent transcription are detected by smFISH. **(A)** Sites of nascent transcription can fluoresce more brightly than single mRNA molecules due to multiple nascent transcripts localized to one gene locus. **(B)** Probes recognizing an *omb* exon generate many small dim spots and a few large bright spots. Right image shows the merge of probe and DAPI fluorescence. The bright spots are associated with nuclei whereas most dim spots are not. **(C)** Probes recognizing an *omb* intron only generate large bright spots that are associated with nuclei. Scale bars = 5 μm. (D) Frequency distribution of intensity for all spots identified in a wing disc probed for *sens* RNAs. Using a threshold of twice the median spot intensity, all single mRNA spots were filtered out, leaving only spots that are associated with transcription sites. The frequency distribution for this class of spots is shown. **(E)** Transcription sites are assigned to cells. For each cell that contains one or more mRNA molecules, it is scored for whether it also has one or more transcription sites. The average fraction of all such cells with a transcription site is shown for each gene. Error bars represent 95% confidence intervals. **(F)** The variance of mRNAs/cell is ratiometrically compared to the mean mRNAs per cells for all genes. This ratio is much larger than one, irrespective of the median mRNA number for binned sub-populations of cells and the gene type. Error bars represent 95% confidence intervals.

Although wing disc cells are diploid, fewer than 15% of nuclei contained more than one transcription site for a given gene. *Drosophila* and other animals have extensive physical pairing of homologous chromosomes in somatic cells (McKee, 2004). Consequently, alleles on paired chromosomes are often spatially juxtaposed (Szabo et al., 2018). Thus, a single transcription spot in a nucleus likely represents transcription from both alleles.

### Transcription occurs in bursts

Transcription sites were counted by applying a threshold that only included spots with at least twice the intensity of a mature mRNA spot (Figure 3D, Figure 3 - figure supplement 1). There was a broad distribution of transcription site intensities, suggesting a large range of nascent RNA numbers that were present on a gene at a given time.

Strikingly, many cells did not have a detectable transcription site even though the cells contained mature mRNAs (Figure 3E). From 50 - 80% of cells had this feature, and it was observed for all genes. This observation is not an artifact of segmentation erroneously assigning mature mRNAs to a cell, since the presence of transcription sites was highly variable before segmentation (Figure 3 - figure supplement 2).

Why do cells with mature mRNAs lack detectable transcription sites? One explanation is that each gene’s promoter is always open, but since transcription is fundamentally stochastic, there would be times when zero or just a few Pol II molecules are Atranscribing the gene. In this scenario, the birth and death of mRNAs can be described as a Poisson process. For simple Poisson processes, the ratio of the variance to the mean number is one. In our case, Poisson-like birth-death of mRNAs would yield a ratio of variance in mRNA number to mean mRNA number to be one (Munsky, Neuert, & van Oudenaarden, 2012; Arjun Raj & van Oudenaarden, 2008). Since mRNA number per cell varied systematically across the wing disc because of Wg and Dpp signaling, we binned cells according to their position in the disc, as had been described earlier (Figure 1G, 2H). Strikingly, the ratio of variance to mean mature mRNA number per cell was between 5 and 10 for all genes, and was also fairly independent of mRNA output (Figure 3F). This indicated that a simple Poisson process could not explain why we failed to detect transcription sites in every cell expressing mRNA.

To determine if our observations were possibly caused by transcription bursting, we invoked a classical two-state model of transcription (Figure 4A). A promoter exists in one of two possible states - ON and OFF. The promoter switches between states at particular rates *k_on_* and *k_off_*. When the promoter is in the ON state, Pol II is permitted to initiate transcription that is subject to a rate constant *k_ini_*. When the promoter is in the OFF state, Pol II is unable to initiate transcription. The model also includes a transcription elongation step, which is assumed to be 100% processive, and whose timescale depends on the gene length and the rate of elongation. The latter is assumed to be 1,100 nucleotides/min, which is a value that has been experimentally determined in *Drosophila* (Ardehali et al., 2009).

**Figure 4.**
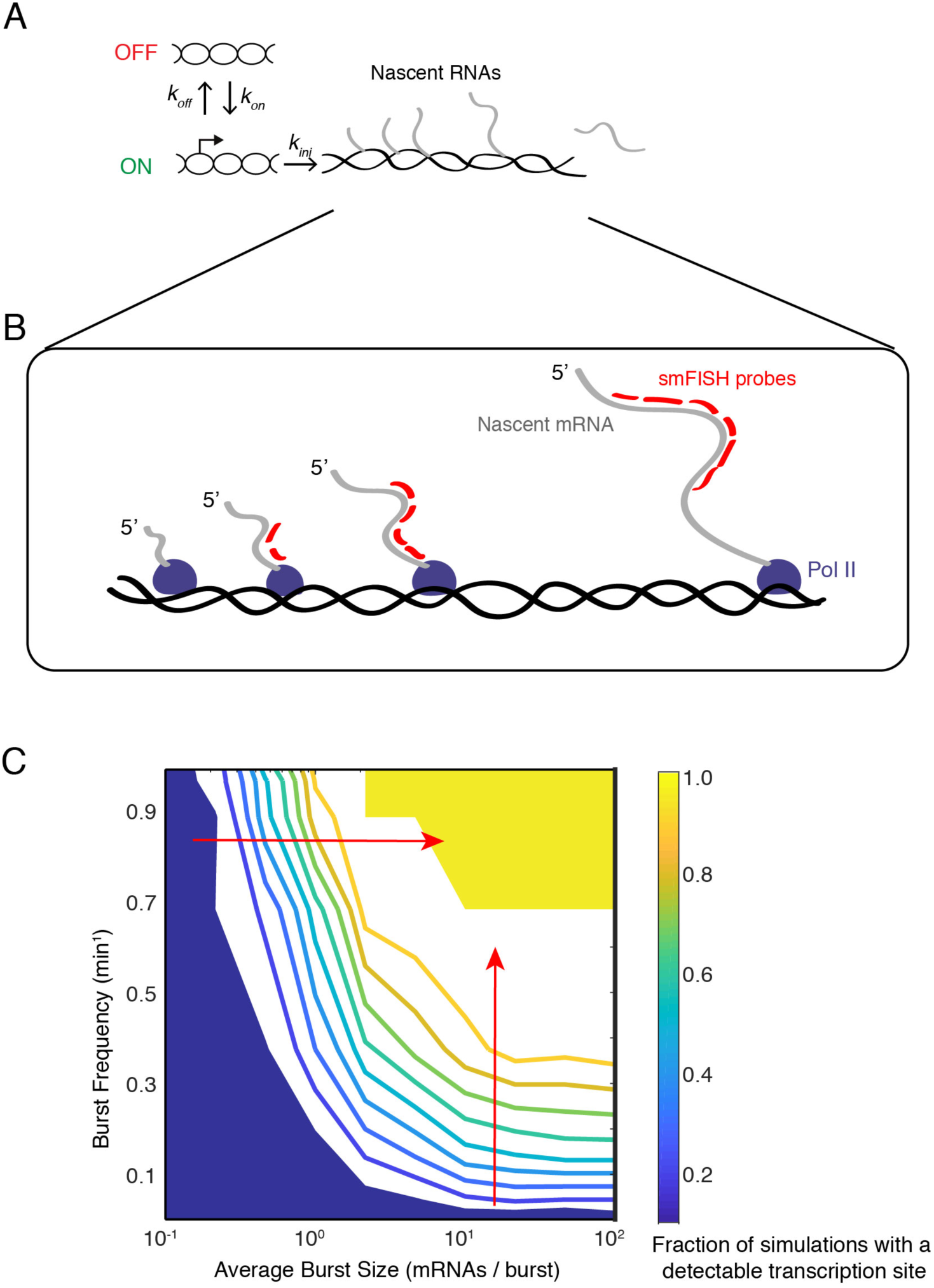
Modeling transcription sites using bursting dynamics. **(A)** Model framework showing the three rate parameters affecting transcription initiation. Two parameters affect the promoter state, while the third parameter only affects how many initiation events occur when the promoter is ON. **(B)** Pol II molecules in elongation mode are distributed along the transcription unit. If Pol II is upstream of the probe binding sites, the nascent transcript will not be detected. If Pol II is downstream, the nascent transcript will be detected as 100% signal. If Pol II is transcribing within the binding sites, the nascent transcript will be detected as a partial signal. These three different scenarios are all found in model simulations. For example in the simulation result shown here, four Pol II’s are situated such that a total of 12 virtual probe binding sites are present. Since each mRNA has 6 binding sites, it means that this simulated transcription site has 12/6 or 2 units of normalized signal. Applying our filter cutoff for identifying a transcription site as 2 or more units, this simulated site would be scored as a positive. **(C)** The distribution of normalized signal intensity for 1000 transcription site simulations. Shown are two distributions from simulations with different initiation rate parameters. Those simulations that result in signal intensities of 2 or more units are classified as detectable transcription sites. **(D)** The phase diagram of transcription site detection in the model. When burst size increases at low burst frequency, the likelihood of detecting a transcription site remains fairly constant. When burst size increases at high burst frequency (horizontal red arrow), the likelihood of detecting a transcription site is ultrasensitive to burst size. Likewise, when burst frequency increases at low burst size, the likelihood of detecting a transcription site remains fairly constant. When burst frequency increases at high burst size (vertical red arrow), the likelihood of detecting a transcription site is ultrasensitive to burst size. There are many values of burst size and frequency that could theoretically explain the observed transcription site frequencies that range from 0.2 to 0.5.

In the model, transcriptional bursts have a characteristic size (number of transcripts per burst) and frequency (rate at which bursts occur). The average burst size is defined as *k_ini_* / *k_off_*, whereas the average burst frequency is defined as (*k_on_^-1^* + *k_off_^-1^*)^-1^. We systematically and independently varied the parameters *k_on_, k_off_,* and *k_ini_* to tune the frequency and size of virtual bursts. For each parameter set, we ran 1,000 simulations of the master equation. To capture the stochastic nature of gene expression, most reactions in the model were treated as probabilistic events, with the exception of transcript elongation time. Therefore simulations with identical parameter values nevertheless gave variable output.

To better relate the results of model simulations to experimental data, we performed the following treatments. First, we randomly paired two independent simulations to mimic the total transcription site activity of paired alleles within a nucleus. Second, we transformed the two simulations’ output to capture the heterogeneity in fluorescence signal intensity at a transcription site. The intensity of each site depends on how many binding sites for probe are present in all nascent transcripts at the site (Figure 4B). This varies with the number of elongating Pol II molecules on the gene and the fraction of molecules that are elongating within or downstream of the region complementary to the probe set. Since this variable is highly dependent upon the position of the complementary region relative to the transcription start and stop sites, we adjusted model conditions to match each particular gene and its region of probe set complementarity. We used these constraints to estimate transcription site intensities from 1,000 pairs of simulations for each parameter set.

When a simulated transcription site intensity fell below the threshold of twice the number of probe binding sites per mRNA, we counted that simulation as having no “detectable” transcription site. This mimicked the threshold that was applied to the experimental data for identifying a transcription site. We then asked what combination of burst size and frequency could theoretically account for the observed frequency of finding cells with a transcription site (this ranged from 20 to 50% of cells). A phase diagram revealed that a broad range of burst sizes and frequencies would explain our observations (Figure 4C). Therefore, according to our model results, tuning burst frequency and/or size can produce a variable likelihood of detecting a transcription site by smFISH.

### Burst frequency is regulated by Dpp and Wg

We quantified the frequency of detecting a transcription site as a function of cell position within the wing pouch (Figure 5A,B). This frequency varied across the disc in a manner that was gene-specific. Strikingly, for all genes, the spatial distributions of transcription site frequency strongly paralleled the mRNA number per cell (compare Figure 5A,B and Figures 1G, 2H). We further examined the relationship between mRNA number per cell and transcription site frequency (Figure 5C,D). Indeed, average mRNA number per cell and the probability of detecting transcription sites in cells were strongly correlated with one another. Remarkably, the slopes of linear fits for three Dpp-responsive genes, *brk*, *omb*, and *salm*, were not significantly different from one another (Figure 5E). The slope for *dad* was similar to *brk* and *omb* but smaller than for *salm*. The slope for *sens* was smaller still. The linear correlation between frequency of transcription site detection and mRNA number confirms that gene regulation by Dpp and Wg is primarily determined through control of transcription initiation.

**Figure 5.**
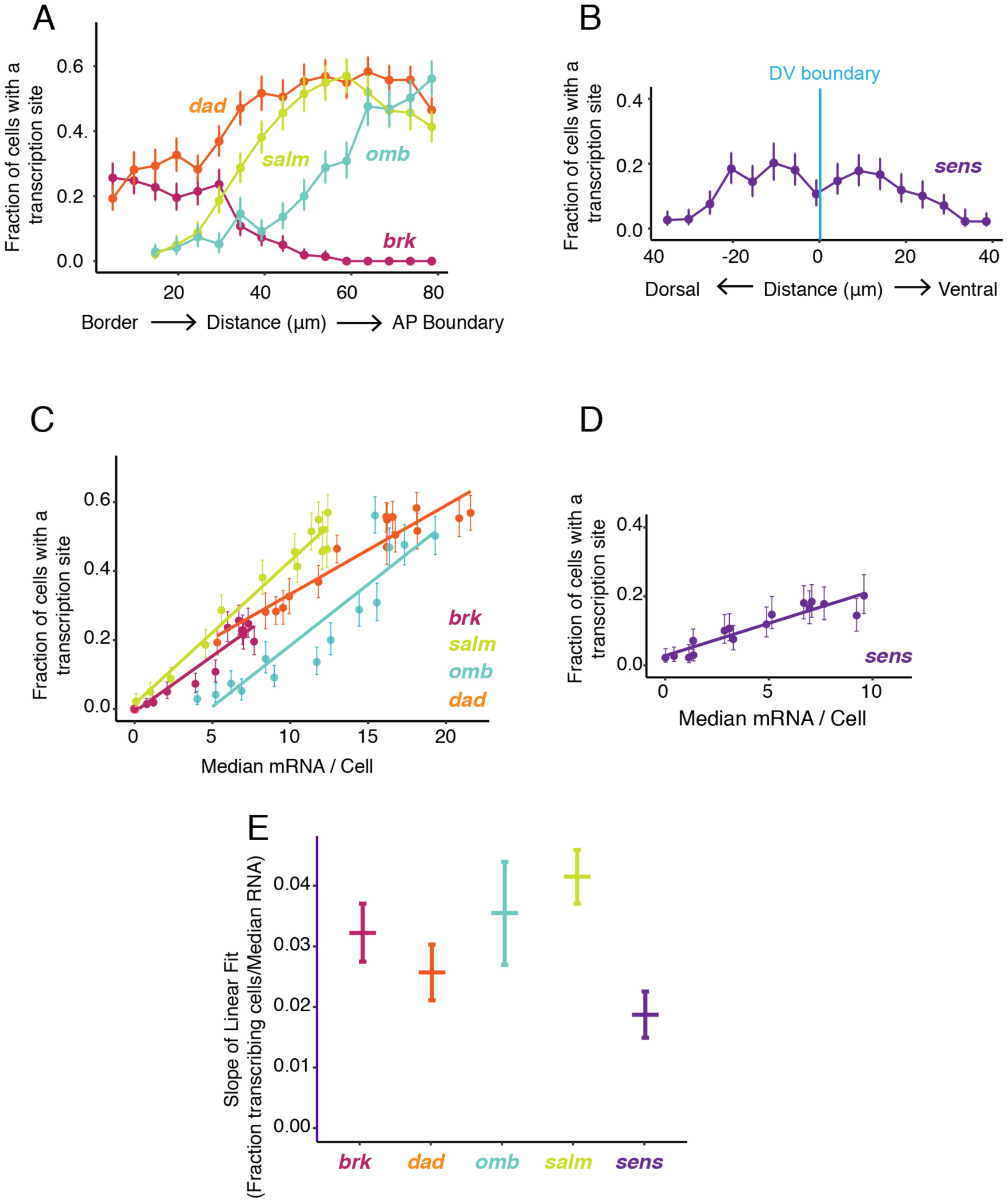
Transcription site detection correlates with mRNA number. **(A,B)** The probability of detecting a cell with a transcription site varies with the cell’s location relative to the source of morphogen. Error bars are 95% confidnece intervals. **(A)** Cells are binned according to their distance from the pouch border, and the fraction of cells in each bin with a transcription site are shown for each Dpp-responsive gene. **(B)** Cells are binned according to their distance from the DV boundary, and the fraction of cells in each bin with a transcription site is shown for the *sens* gene. **(C,D)** The probability of detecting a cell with a transcription site varies linearly with the number of mRNA molecules in the cell. Fitted lines are from linear regression. Error bars are 95% confidence intervals. **(C)** Cells are binned according to the number of mRNAs they contain, and the fraction of cells in each bin with a transcription site are shown for each Dpp-responsive gene. **(D)** Cells are binned according to the number of mRNAs they contain, and the fraction of cells in each bin with a transcription site is shown for the *sens* gene. **(E)** Linear regression analysis was performed on samples from C and D, shown is the slope with a parametric 95% confidence interval.

The likelihood of detecting a transcription site increases because either the promoter is spending more total time in the ON state or more RNAs are being transcribed while in the ON state. These properties are affected by burst size and burst frequency in different ways. We sought to determine whether burst size or frequency was being regulated. We did so by estimating the number of nascent RNAs at each transcription site, which was quantified as a multiple of the median pixel intensity of mature RNA spots (Figure 3 - figure supplement 1). The average number of nascent RNAs per transcription site did not significantly vary between cells that were receiving different levels of Dpp and Wg signal (Figure 6A,B). This was observed for all genes. Moreover, the average number of nascent RNAs per transcription site was also independent of the likelihood that transcription was occurring in a cell (Figure 6C). Therefore, the propensity for a cell to generate nascent transcripts does not correlate with the number of nascent transcripts.

**Figure 6.**
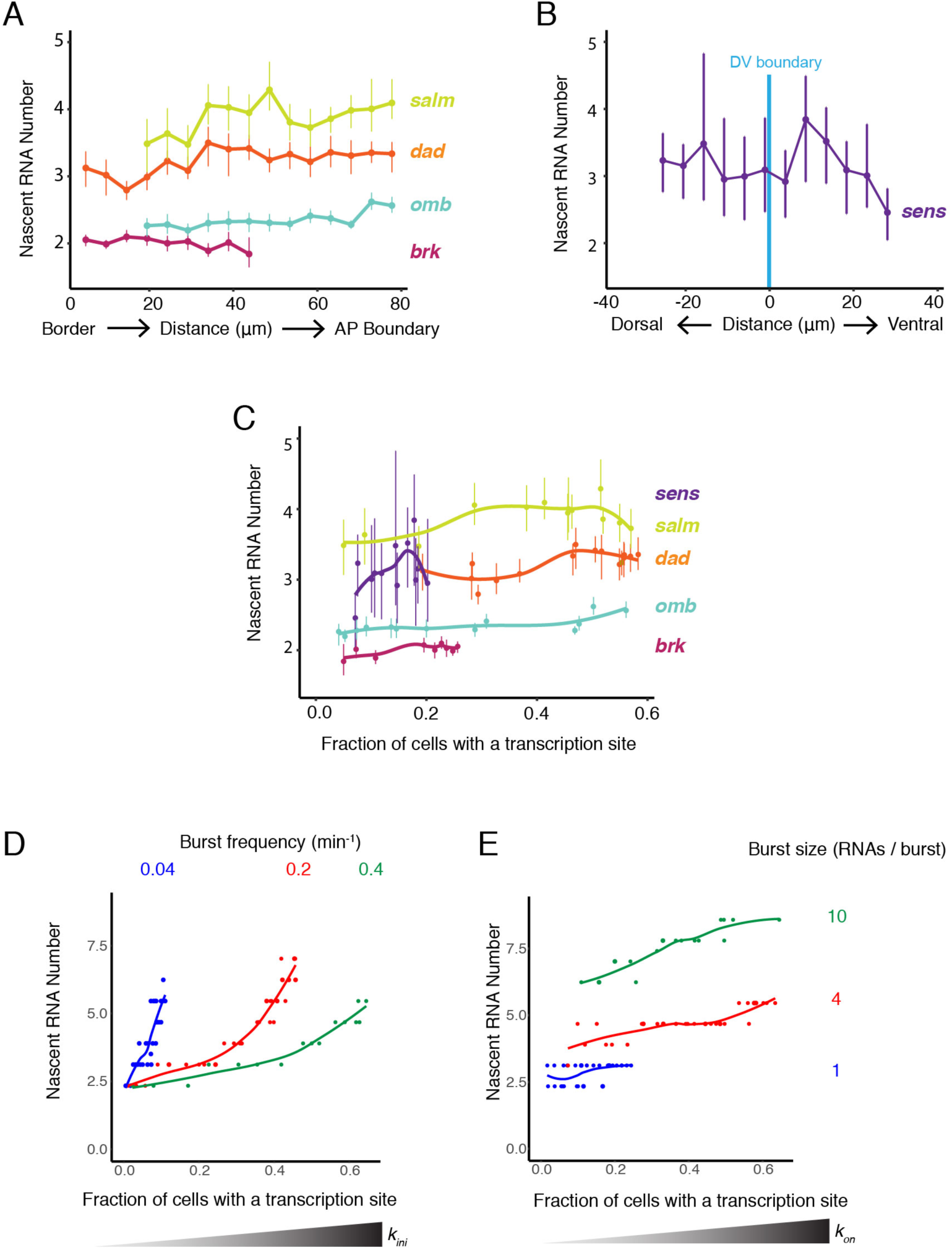
Burst frequency is regulated by Dpp and Wg. **(A,B)** The average number of nascent RNAs in a transcription site does not vary with the cell’s location relative to the source of morphogen. Error bars are bootstrapped 95% confidence intervals. (A) Cells are binned according to their distance from the pouch border, and the average number of nascent RNAs per site in each bin are shown for each Dpp-responsive gene. (B) Cells are binned according to their distance from the DV boundary, and the average number of nascent RNAs per site in each bin is shown for the sens gene. (C) The average number of nascent RNAs in a transcription site does not vary with the probability of detecting a cell with a transcription site. Error bars are 95% confidence intervals. (D,E) Modeling the relationship between average number of nascent RNAs in a transcription site and the probability of detecting a site for the dad gene. (D) Simulations are performed where the rate parameter kini has been systematically varied so that burst size is variable. Resulting values for nascent RNA number and fraction of cells with a site are shown. Each datapoint is the average of 1,000 simulations. Simulations are repeated for three different values of kon to specifically set the burst frequency to 0.04, 0.2 and 0.4 min-1. Simulations are performed where the rate parameter kon has been systematically varied so that burst frequency is variable. Resulting values for nascent RNA number and fraction of cells with a site are shown. Each datapoint is the average of 1,000 simulations. Simulations are repeated for three different values of kini to specifically set the burst size to 1, 4 and 20.

To understand the relationship between these observed features, we turned to the modeling framework. We first considered whether modulation of transcription burst size by Wg and Dpp could explain our observations. We modulated burst size by systematically varying the *k_ini_* parameter, and from simulations, then calculated the number of nascent RNAs per transcription site and the transcription site detection frequency. There was a positive correlation between nascent RNA number in a transcription site and the probability of detecting a transcription site (Figure 6D and Figure 6 - figure supplement 1A). This was observed across a wide range of fixed burst frequencies. When nascent RNA number was 3 or higher, the correlation with transcription site frequency was strongest. Moreover, when the probability of a transcription site was very low, nascent RNA number converged to a common value irrespective of burst frequency. None of these model predictions were observed in the experimental results with the target genes (Figure 6C). It suggests that transcription burst size is not strongly regulated by Dpp and Wg.

We then modulated burst frequency in the model by systematically varying *k_on_*, and calculated the number of nascent RNAs per transcription site and the transcription site frequency. There was little change in nascent RNA number as transcription site frequency changed, even across a wide range of fixed burst sizes (Figure 6E and Figure 6 - figure supplement 1B). Interestingly, the burst size appeared to determine what nascent RNA number value was held at a constant. Moreover, there was no convergence of nascent RNA number when the probability of a transcription site was very low, irrespective of burst size. All of these model predictions agree well with the experimental results (Figure 6C). This suggests that Dpp and Wg regulation of genes in the wing disc primarily occurs by modulation of transcriptional burst frequency.

## DISCUSSION

Morphogens elicit different transcriptional outputs from target genes, depending on local concentration of the morphogen. The targets of Dpp signaling in the wing offer a well-studied example of this concept. Transcription of the gene *brk* is directly regulated by Mothers-against-dpp (Mad), the effector of Dpp (Minami, Kinoshita, Kamoshida, Tanimoto, & Tabata, 1999; Moser & Campbell, 2005). In complex with Medea and Schnurri, Mad represses *brk* transcription (Cai & Laughon, 2009). This generates a gradient of Brk protein expression that is inverted to the Dpp gradient. In turn, the level of Brk protein is instrumental in repressing the expression of genes that are induced by Dpp, including *omb* and *salm* (Campbell & Tomlinson, 1999). Thus, opposing gradients of activation and repression define the expression domains of *omb* and *salm*. Since *omb* is less sensitive to Brk repression than *salm*, its expression domain is broader than that of *salm*. Transcription of *salm* is directly activated by Dpp, but in this case, Mad and Medea without Schnurri activate *salm* transcription (Moser & Campbell, 2005). Curiously, *omb* transcription does not directly depend on Dpp signaling, and its transcriptional activation is brought about by unknown factors (Sivasankaran, Vigano, Muller, Affolter, & Basler, 2000).

Given the diverse mechanisms by which genes such as *omb*, *brk*, and *salm* are regulated, it is illuminating that the frequency of transcription bursting is the regulated step for all of these genes. Burst size appears to be independent of Dpp signaling for these genes. If our two-state model for initiation is accurate, then *k_on_* is the most likely step that is being regulated directly and indirectly by Dpp. This is because *k_on_* specifically affects burst frequency and not size whereas *k_off_* affects both frequency and size. If *k_on_* is the kinetic step under regulation for all of these genes, how is it rendered rate-limiting given such diverse enhancer architectures and transcription factor inputs? It has been found that burst frequency correlates with enhancer strength and enhancer-promoter contact, suggesting that *k_on_* is potentiated by enhancer-promoter contact and is mediated by transcription factor binding to DNA (Bartman et al., 2016; Bothma et al., 2014; Chen et al., 2019; Fukaya et al., 2016; Larsson et al., 2019).

In spite of this universal regulation of burst frequency by Dpp, there are other factors that also help determine the expression domains of these genes. For example, *salm* and *omb* are expressed in nested domains; cells close to the source of Dpp contain mRNAs for both genes while lateral cells more distant from the source contain predominantly *omb* mRNAs (Figure 2H). Although burst frequency control might solely dictate these differences, it appears not to be the case. The likelihood a cell is transcribing either gene does not strictly correlate with the breadth of their expression domains. Lateral cells that predominantly contain *omb* mRNAs nevertheless are more likely to be transcribing *salm* than *omb* (Figure 5A). Looking at the relationship between transcription likelihood and mRNA number (Figure 5C), the relationship for both genes is linear with similar slopes. But for *omb*, the curve is shifted such that more mRNAs are found in cells that have a lower likelihood to be undergoing transcription. This shift is not due to a greater burst size of *omb* transcription (Figure 6C). Rather, the simplest interpretation is that the half-life of *omb* mRNA is greater than the half-life for *salm* mRNA, so *omb* mRNAs are more readily detected in cells between transcription bursts. Thus, the broader domain of *omb* expression might be accounted for by mRNA stability.

This conclusion is at odds with previous studies of *omb* regulation that used an enhancer trap reporter for *omb* expression. There, the reporter was expressed in a broad domain within the wing. However, this reporter expressed lacZ in response to *cis*-regulatory elements near the most distal promoter driving a minor species of *omb* transcript (Mayer, Diegelmann, Abassi, Eichinger, & Pflugfelder, 2013). There is a more proximal promoter 13 kb that appears to be the major site of transcription initiation for *omb* (Flybase).

Our results also challenge the view that *salm* and *omb* expression domains have sharp boundaries due to transcription thresholds set by Brk and Dpp. We find that *omb* and *salm* mRNA numbers per cell drop gradually with distance from the source of Dpp (Figure 2H). As well, their gradients in mRNA number are inversely correlated with the gradient in *brk* mRNA number. *Salm* has relatively constant mRNA number in cells near the AP boundary, and those numbers gradually diminish in cells located more laterally. A similar pattern is seen with *omb*, except the domain with constant *omb* mRNA number is smaller than for *salm*. However, the *salm* and *omb* enhancer trap reporters as well as anti-Salm immunohistochemistry have reported expression domains with sharp boundaries. Possibly, the discrepancy hints at some threshold of mRNA expression below which protein output drops sharply. It is also possible that the previously characterized expression domains for *salm* and *omb* were distorted by non-linear detection of antibodies that recognize Salm and the protein product of lacZ, β-galactosidase.

## MATERIALS AND METHODS

### Drosophila genetics

All *Drosophila* were raised at room temperature and grown on standard molasses-cornmeal food. The *sfGFP*-*sens* transgenic line was used as described in (Cassidy et al., 2013). Experiments were performed on *dad-GFP* and *brk-GFP* transgenes obtained from Bloomington Drosophila Stock Center (stocks 81273 and 38629, respectively). For all transgenic experiments, smFISH was performed on homozygous individuals. Experiments were performed on endogenous *omb* and *salm* in *w^1118^* individuals.

### smFISH Probe Design and Preparation

smFISH oligonucleotide probes were designed using Stellaris Probe Designer (Biosearch Technologies). Probes sets contain between 45 and 48 non-overlapping 20-nucleotide oligos. A full list of all probe sets is provided in Supplementary Table 1. Anti-GFP probes were prepared by conjugating NHS-ester ATTO 633 dye (Sigma 01464) to the 3’ end of each oligonucleotide. Anti-Sens probes were prepared by conjugating NHS-ester ATTO 565 dye (Sigma 72464) to the 3’ end of each oligonucleotide. These oligos bear a mdC(TEG-Amino) 3ʹ modification to allow conjugation, and were obtained from Biosearch Technologies. Conjugation and purification was performed as described (S. C. Little & Gregor, 2018). All other probe sets were prepared using the enzymatic conjugation protocol as described (Gaspar, Wippich, & Ephrussi, 2017). Briefly, amino-11-ddUTP (Lumiprobe) was conjugated to NHS-ester ATTO 633. Terminal deoxynucleotidyl transferase (New England Biolabs) was then used to conjugate ATTO 633-ddUTP to the 3’ ends of oligonucleotides that had been purchased from IDT. After enzymatic conjugation, oligos were purified from free ATTO 633-ddUTP using G-25 spin columns (GE Illustra) according to manufacturer’s instructions. Final concentration of oligonucleotide was 33 µM in water. Probes were stored at −20°C, protected from light, until use.

### smFISH

Wing discs were dissected from wandering 3^rd^ instar larva in cold phosphate buffered saline (PBS) and immediately fixed in 0.1% (w/v) paraformaldehyde / PBS for 15 minutes at room temperature. Discs were then fixed for 30 minutes in methanol at room temperature. Discs were transferred to hybridization buffer (10% w/v dextran sulfate, 4X SSC, 0.01% w/v salmon sperm ssDNA (Invitrogen 15632), 1% v/v vanadyl ribonucleoside (NEB S14025), 0.2mg/mL BSA, 0.1% v/v Tween-20). Oligo probes were added to a 1.5 µM final concentration in the hybridization buffer, and hybridization was performed for 1 hour at 62° C. After hybridization, discs were washed once for 5 minutes at 62 ° C in wash buffer (4X SSC, 0.1% v/v Tween-20). Discs were then incubated with 2.5 μg/mL 4ʹ,6-diamidino-2-phenylindole (DAPI) (Invitrogen) in PBS + 0.1% Tween-20 for 5 minutes at room temperature. Discs were washed with PBS + 0.1% Tween-20 and transferred to Vectashield (Vector Labs) for mounting. Discs were mounted in 15 μl of Vectashield on glass microscope slides using an 18 × 18 mm No. 1 coverslip (Zeiss). For eye imaginal discs, discs were dissected from late 3^rd^ instar larva in cold PBS with brain and mouth hooks attached, then smFISH was performed as described. Immediately prior to mounting, brain and mouth hooks were removed from eye discs and discarded.

### Actinomycin D Treatment

Wing discs were dissected in room temperature Graces’ Insect Medium (Sigma 69771) supplemented with 1X Pen-Strep (Gibco 15140-122) and 5 mM Bis-Tris (Sigma B4429). Half of the total dissected discs were transferred to 24-well tissue culture dishes containing this prepared media + 5 ∝g/mL Actinomycin D, and half were transferred to untreated controls containing culture media + 1:1000 (v/v) DMSO. Discs were incubated with gentle shaking for 30 minutes at room temperature, protected from light, before being washed with fresh culture media, and 1X PBS. SmFISH was then performed as described.

### Microscopy

12-bit 3D image stacks were collected with x-y pixel size of 76 nm and z-intervals of 340 nm on a Leica SP8 scanning confocal microscope, using a pinhole size of 1 Airy unit and a 63X oil immersion (NA 1.4) objective. DAPI, ATTO 565, and ATTO 633 were excited by the 405, 555, and 630 nm lasers, respectively. ATTO dye fluorescence was collected using a HyD detector on photon counting mode and a scanning speed of 200 Hz, with 16X line accumulation. DAPI fluorescence was collected using PMT detector using 8X line averaging.

### Image Processing

Raw smFISH images were processed using a custom Matlab pipeline with no prior preprocessing. Our pipeline is available at github.com/bakkerra/smfish_pipeline. The pipeline consists of several modules.

### Selection of mRNA Segmentation Threshold

RNA segmentation is performed by applying a threshold value to an smFISH image, and transforming all pixels above the threshold (‘objects’) to white and pixels below this threshold (‘background’) to black. To robustly identify RNAs, it is therefore important to select a threshold where real RNA fluorescent spots are above the threshold, and background fluorescence is below the threshold. Using this threshold method, we classify an object in each 2D image when white components have a connectivity of 8 pixels or more. When the number of objects in an image stack is counted across a range of segmentation thresholds, the number of objects reaches an inflection point and plateaus at a threshold approximately equal to the level of fluorescence that separates real RNA spots from background (see Figure 1 - figure supplement 1C).

We manually identified and labeled 347 RNA spots from sub-regions of four independent image stacks, and found that when a threshold is selected within the plateau after the inflection point, the number of objects identified is no more than +/− 5% different than the ground truth manual curation. Furthermore, the centroids of identified objects have an average displacement of only 2 pixels from the manually identified centroids. Therefore, this plateau is an appropriate threshold for accurate segmentation of RNA spots.

We reasoned that the position of inflection might vary from sample to sample. Therefore, for each image stack, a range of thresholds is tested, and a threshold is selected within the plateau to collect the segmented data. As a result, each image stack has the potential for a unique threshold, allowing robust segmentation of spots despite variation in raw fluorescence between image stacks. In practice, replicates from the same experiment captured in the same imaging session did not require thresholds for segmentation more than 15 fluorescent units apart. If image stacks did not show an identifiable inflection point and plateau, the signal-to-background of that sample was determined to be insufficient and it was not used for analysis. The smFISH protocol and imaging is robust enough that in our hands, this occurs in less than 10% of image stacks collected. Once a threshold is selected, the following properties of each object are recorded: x-y centroid position, z-plane, and a list of the connected pixels.

### Connecting Segmented Objects into mRNA Spots

Diffraction-limited fluorescent spots captured with the 63X objective at 633 nm wavelength are estimated to be approximately 600 nm in diameter. This corresponds to a diameter of 8 x-y pixels in our images (Lipson, 1995). As each z-plane is 340 nm in depth, it is assumed that genuine diffraction-limited RNA spots will appear in 2 or 3 consecutive z-planes, depending on the spot’s position along the z-axis.

Therefore, candidate RNA spots must satisfy two criteria in order to be counted:

1. Candidate must have a corresponding object centroid at least one neighboring z-plane within a diffraction limited radius of 4 pixels.
2. Candidate must be larger (contain more pixels) than corresponding objects in neighboring z-planes. This criterion prevents RNA fluorescence spots from being counted in multiple z-planes.

A candidate that satisfies these criteria is recorded as an mRNA spot, and only the largest 2-D object is recorded.

The pipeline shows the FISH images overlaid with markers indicating recorded spots so that each image stack can be manually inspected for any significant errors or inconsistencies. The most common problem detected at this stage resulted from images taken of discs that were “drifting,” or moving significantly between z-slices, which can cause a large number of identified spots to be filtered out during processing for not meeting criterion 1. Excessive bleaching across the z-stack can also cause clear inconsistencies. In this study, such problems were rare enough that any sample experiencing these problems was considered to have failed quality control and was simply not included for further analysis.

Intensity measurements are recorded from a circle of pixels of radius 4 about the centroid of each recorded RNA spot. By keeping the area of each intensity measurement fixed, we uncouple user-generated variation in selection of segmentation thresholds from spot intensity measurements. A 2D circle was used instead of a 3D sphere to extract intensity measurements because the spots only appear in 2 or 3 z-planes. This makes their 3D geometry variable from spot to spot, and they cannot be consistently described using a sphere or ellipse. **Segmentation of Transcription Sites:** In our images, transcription sites tend to contain pixels that are many times brighter than mature RNA spots. As a result, the brightest transcription sites are frequently misidentified during segmentation of mature RNA spots because the second criterion for spot identification only records the largest object within the diffraction limit in z. For transcription sites, this is not always the brightest plane. Therefore, we segment transcription sites independent of mature RNAs using a higher threshold. The objective in threshold selection for transcription sites is to select one that includes objects with a total fluorescence intensity of twice the average mature RNA, and excludes mature RNA spots. We define the “average” intensity for a spot containing a single mRNA to be the median of the distribution of all identified mature RNA spot objects. We empirically determined that merely doubling the threshold for segmentation does not achieve this, because mature mRNAs may contain a few pixels above the threshold, enough to still be identified as objects and included in analysis. Therefore, we use a threshold calculated by multiplying the median mature RNA intensity by a factor of 2.5 (Figure 3 -figure supplement 1A).

To test the accuracy of this segmentation procedure, we manually inspected three particularly RNA-dense regions in independent images where automated segmentation found a total of 103 transcription sites and 4,066 mature RNAs. We determined that only 7 of 4,066 mature RNAs were misidentified as transcription sites, and found no examples of transcription sites that had been missed by automated segmentation.

After identification, object intensity measurements are recorded from a circle of pixels of radius 4 (the diffraction limit) about the centroid of each identified transcription site (Figure 3 -figure supplement 1B). The average transcription site threshold selected for replicates in a dataset show no correlation with the average intensity of transcription sites in that dataset (Figure 3 -figure supplement 1D). Therefore, the differences in transcription site intensity between genes cannot be explained merely by differences in threshold selection or variability in image fluorescence between datasets.

### Estimation of Nascent RNA Number per Transcription Site

The intensity measurement of each identified transcription site in an image stack is divided by the median intensity of identified mature RNAs in that sample (Figure 3 -figure supplement 1C). This serves two purposes. First, it serves to normalize these measurements within each sample so transcription site intensity measurements can be pooled across replicates without the effects of image-to-image variability in fluorescence. Secondly, each transcription site object is presumed to be the sum of intensities of multiple nascent RNA molecules elongating at the transcription site. By dividing each transcription site intensity by the average intensity of a single RNA, we obtain an estimate of the number of nascent RNAs present at the transcription site. Because some transcripts are partially elongated, this number cannot be completely accurate, and we attempt to compensate for this in our computational model when interpreting results.

### Nuclei Segmentation

DAPI fluorescence images are output as labeled 16-bit images, where each nuclear object corresponds to a ‘level’ in the 16-bit image. These images are input to a nuclei segmentation pipeline, which flattens the images to white nuclei objects and black background. Nuclei images are segmented in 2D using the NucleAIzer platform maskRCNN Network, trained as described in (Hollandi, 2019) This is available online at nucleaizer.org. We trained the neural network with an expected nuclear radius of 32 pixels (Figure 1 - figure supplement 2D). To ascertain the accuracy of segmentation, we compared results to manually labeled nuclei in four randomly selected disc images. The automated method identified at least 85% of nuclei objects identified manually for each image.

The segmented black and white images are then processed using a custom Matlab script in order to join overlapping 2D objects into 3D. Each nucleus object in each z-slice is assigned an identity index. For each object in the first z-slice, the object with the highest number of overlapping pixels in the next z-slice is identified, and this object’s identity index is altered to be identical to its overlap partner. This proceeds through the entire z-stack of images, creating objects that resemble ‘pancake stacks’ of linked 2D objects in 3D (Figure 1 - figure supplement 2E). The 3D-centroid and list of included pixels of these new objects is then recorded. Objects not incorporated into a 3-D object are disregarded.

### Generation of Voronoi Diagrams

A 3D Voronoi tessellation divides a geometric volume into spatial regions with boundaries equidistant from a set of points (Voronoi, 1908). We use this method to assign RNA objects to the nearest nuclear centroid. The set of segmented 3D nuclear centroids are used to divide the z-stack of images into a 3-D Voronoi tesselation using a polytope bounded Voronoi diagram available for Matlab, which uses the DeLaunay triangulation to calculate the Voronoi diagram (Park, 2020). The result of this tesselation is a list of 3-D vertices of each Voronoi ‘cell’ in space, which is recorded along with the associated nuclear centroid (Figure 1 - figure supplement 2F).

### Assignment of RNA to nearest nuclei

Mature mRNAs and transcription spots located within a Voronoi spatial cell are assigned to that particular cell. To assign spot objects to cells, a 3D convex hull of the each Voronoi cell is constructed from the vertices data for that cell. An entire set of image points, either the mRNA or transcription spot centroids, are tested to determine whether they fall inside or outside of each hull (Figure 1 - figure supplement 2E). This is performed using a Matlab function called inhull, which uses dot products to shorten calculation times (D’Errico, 2020). Spots that fall inside a given cell’s Voronoi hull are assigned to that cell’s nuclear centroid, and the number of assigned spots, as well as their centroid and z-plane information are recorded. This is then repeated for every Voronoi cell in the image stack. The final result is a list of cells, their nuclear centroids, the total number of RNA spots assigned, and a list of each assigned spot’s centroids.

## Data Analysis

### Binning of data

Each disc is imaged with the DV boundary located at the y-coordinate midline of the image. Therefore the x-coordinate of the image corresponds to position along the disc’s AP axis, and the y-coordinate corresponds to position along the DV axis. In order to analyze data across developmental axes, each image is divided into spatial bins of 64 pixels each, approximately equal to the diameter of one cell nucleus. RNA spots are assigned to a bin according to the position of their associated nuclear centroid.

### Sample size and replicates

We analyzed image stacks from three independent discs for each experiment. Each image stack contains approximately 1,700 identified nuclei. Therefore, the total sample size is approximately 5,000 cells per experiment. Similar trends in RNA and transcription spots feature are observed in each disc individually, and hence, the analysis is not distorted by artifacts in pooling and cell segmentation (Figure 3 - figure supplement 2).

### Alignment of replicates along developmental axes

While each disc is imaged roughly in the same region, there is not an unambiguous landmark that precisely registers different disc images with one another. To pool data across space as accurately as possible, we register discs to each other based on their mRNA spot distributions over space. For each image data set, the number of RNAs per spatial bin is summed, and the distributions across bins are compared. Bins are then manually registered such that the distribution profiles of the 3 datasets line up with one another (Figure 3 - figure supplement 2A-E). The overlapping bins from the three datasets are then assigned to a pooled bin. Pooling includes the nuclei centroids as well as the transcription and RNA spots. This is repeated for all bins.

### Calculations

*Median mature mRNAs per cell* is calculated from total number of mature mRNA spots for each cell within a spatial bin of pooled data. As the distribution of mRNAs per cell is not normally distributed and has a long tail, we ascertained that the median was a more robust descriptor of the “center” of the distribution than mean.

*Median nascent RNAs per cell* is calculated from normalized intensity measurements for each transcription spot within a spatial bin of pooled data. All nascent RNA spots are included. As the distribution of RNA per cell is not normally distributed and has a long tail, we ascertained that the median was a more robust descriptor of the “center” of the distribution than mean. Because the number of transcription sites varies over space, sample sizes vary for calculating median nascent RNAs per cell. For bins where fewer than 5% of cells contain a transcription site, median nascent RNAs per cell was not calculated, as the sample size was determined to be too small (<15).

*Fraction of cells with a transcription site* is calculated by dividing the number of cells in a pooled spatial bin with at least one transcription site assigned to them by the total number of cells in that spatial bin.

*Fano factor* is calculated for each spatial bin by dividing the variance in the mRNA per cell distribution by the mean mRNA per cell for all cells assigned to that pooled spatial bin.

*Curve and line fitting:* Linear models are produced by unweighted least squares linear regression. LOESS fits are performed using the loess fitter in R, with an optimized span to minimize residuals.

*Statistics* are calculated by bootstrap resampling analysis using the bias-corrected and accelerated method. We resample data within each bin of pooled data and calculate the statistic of interest 10,000 times. The mean value of the statistic and a 95% confidence interval are calculated from these resampled values.

### Stochastic Simulation Model

We model the various steps of gene expression, based on central dogma, as linear first order reactions. To simulate the stochastic nature of reactions, we implement the model as a Markov process using Gillespie’s Stochastic Simulation Algorithm (Gillespie, 1977). Simple Markov processes can be analyzed using a chemical master equation to provide a full probability distribution of states as they evolve through time. The master equation defining our gene expression Markov process is as follows:

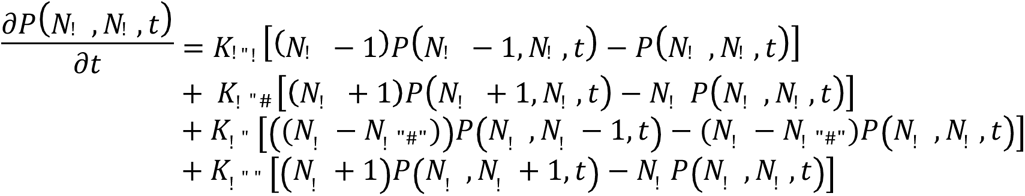

where *N*_!_, *N*_!_, and *t* are defined as the number of RNA molecules present, as the number of transcriptionally active gene copies, and simulation time, respectively. *N*_!“#”_ is defined as the total number of gene copies present, and thus is the maximum number of active gene copies that can exist in the simulation. *K_ini_, k_deg_, k_on_,* and *k_off_* are rate constants defining the rates of transcription initiation, RNA degradation, promoter state switching from off to on, and promoter state switching from on to off, respectively.

As the Markov process gets more complex, the master equation can become too complicated to solve. Gillespie’s Algorithm is a statistically exact method that generates a probability distribution identical to the solution of the corresponding master equation given that a large number of simulations are realized.

A brief description of how the Gillespie simulation produces each probability distribution is as follows:

1. We initialize all simulations to start with no mRNA molecules and promoter state is set to OFF.
2. For each event *i* in the simulation, a total rate *r_tot_* is calculated by summing all *r_i_* reaction rate constants in the model, given the current promoter state and the total number of mRNA molecules present.
3. A time-step τ is generated from an exponential probability distribution with mean 1/*r_tot_*. This τ is the time interval between the current event and the next event.
4. Each event *i* is selected from the list of reaction steps in the model available at that time (promoter switching, transcription initiation, mRNA decay). The probability a reaction step is selected is equal to *r_i_ / r_tot_*. An event is selected at random given these probabilities. For each event, the following actions are taken:

- Promoter switches to ON: Promoter is now in ON state, transcription initiation is now included in r_tot_,
- Promoter switches to OFF: Promoter is now in OFF state, transcription initiation is no longer included in r_tot_.
- Transcription Initiation: Number of mature mRNA molecules is increased by 1.
- RNA degradation: Number of mature mRNA molecules is decreased by 1.
5. Simulation time is updated as *t* + τ where *t* is the total time elapsed in the simulation.

Each simulation is run for 10,000 iterative events to approximate steady-state conditions, at the end of which the number of mRNA molecules present in the simulation is recorded. Independent simulations are then randomly paired to mimic the two alleles within a cell, and the sum of mRNA numbers is recorded as the mRNA output per cell. A minimum of 1,000 simulation pairs are generated for each set of rate parameter values.

The RNA decay parameter *k_deg_* is fixed at 0.04/min for all simulations, as this rate had been experimentally determined for *sens* mRNA (Giri et al., 2019). The transcriptional rate parameters are varied in accordance with the specific hypothesis being tested. We constrain them loosely to be within an order of magnitude of reported values for these rates from the literature (Milo, Jorgensen, Moran, Weber, & Springer, 2010). We also constrain these rates so as to produce steady state mRNA numbers similar to experimental data.
- *k_ini_* is varied from 0.2 to 60 /min
- *k_on_* is varied from 0.008 to 38/min
- *k_off_* is varied from 0.016 to 20/min

To perform a parameter sweep, we vary the relevant parameter across the defined range. Each rate parameter value in the sweep is used to make 1,000 paired simulations as described above.

### Nascent Transcripts

Thus far we have described how model simulations generate *in silico* data for mature mRNA numbers. We also use the same simulations to approximate the number of nascent RNAs per gene. After 10,000 iterative events are completed in a simulation, the number of nascent RNAs is counted. A single nascent RNA is counted if a single transcription initiation event has occurred within an interval of time (τ_elong_) equal to the time it is estimated that RNA polymerase takes to elongate from the binding site for the 5’-most oligo probe to the 3’ end of the RNA. To calculate τ_elong_ for each gene, we divide the number of nucleotides from 5’ probe-binding site to 3’ end by the transcription elongation rate. This rate is assumed to be 1,100 nucleotides/min, as experimentally determined (Ardehali et al., 2009).

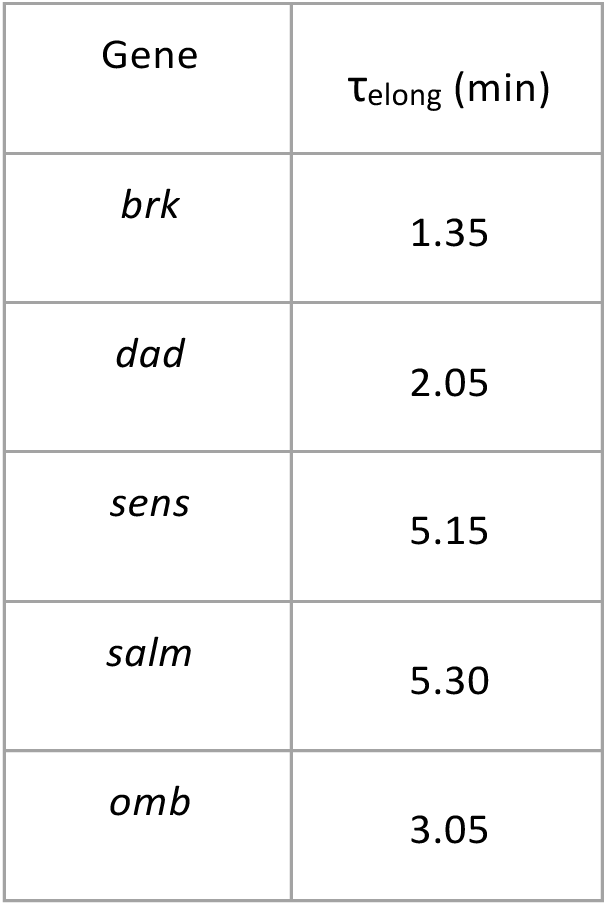

We weight the count of nascent RNAs in a simulation to mimic the fluorescence output from these nascent RNAs if they are hybridized to probes. We define τ_probe_ to be the time interval for RNA polymerase to elongate from the 5’-most probe-binding site to the 3’-most probe-binding site. If a nascent RNA had been initiated in a time less than τ_probe_, then we weight the counting of that nascent RNA as 0.5 rather than 1. We do this because the probe-binding region of the nascent RNA is partially transcribed at this point. For simplicity, the exact locations of probes and RNA polymerase are not taken into account to calculate the weighting, and instead we assign the overall probability of fluorescence for an ensemble of such partially transcribed RNAs. If a nascent RNA had been initiated in a time greater than or equal to τ_probe_ and less than τ_elong_, then we weight the counting of that nascent RNA as 1. These RNAs are assumed to produce 100% of the fluorescence of a mature RNA spot, since all probe-binding sites are transcribed at this point.

We randomly pair two simulations and sum the number of weighted nascent transcripts. This mimics the experimental conditions where the two gene alleles are physically paired and thus their nascent RNAs are co-localized in space. We collate 1,000 paired simulations for each parameter set and calculate the following statistics:

*Fraction of virtual cells with a transcription site* is calculated by counting how many paired simulations have a total number of weighted nascent RNAs of 2.0 or more. This is done in order to be consistent with the limitations of the experimental data; only nuclear spots with fluorescence greater or equal to 2 mature mRNA spots were called as transcription sites. When this number of paired simulations is divided by the total of 1,000 paired simulations, it is the fraction of virtual cells with a transcription site.

*Median number of nascent RNAs per virtual cell* is calculated from those paired simulations with a total number of weighted nascent RNAs of 2.0 or more.

## ACKNOWLEDGEMENTS

Fly stocks from Hugo Bellen and the Bloomington Drosophila Stock Center are gratefully appreciated. We thank Jessica Hornick and the Biological Imaging Facility for help with imaging and the Keck Facility at Northwestern for help with probe purification. We are very grateful to Shawn Little and Thomas Gregor for hosting R.B. at Princeton and their invaluable advice on adapting the smFISH method to imaginal discs. We also thank Arjun Raj and Brian Munsky for key suggestions on experimental and analytical development. Financial support was provided from the NIH (T32CA080621, R.B.) (R35GM118144, R.W.C.), NSF (1764421, M.M and R.W.C.), and the Simons Foundation (597491, M.M. and R.W.C.). M.M. is a Simons Foundation Investigator.

## AUTHOR CONTRIBUTIONS

The experimental work was conceived by R.B. and R.W.C, and R.B. performed all experiments. All data analysis and modeling were performed by R.B. under the supervision of M.M. and R.W.C. The manuscript was written by R.B. and R.W.C. with input from all authors.

---

**Figure 1 - figure supplement 1.**
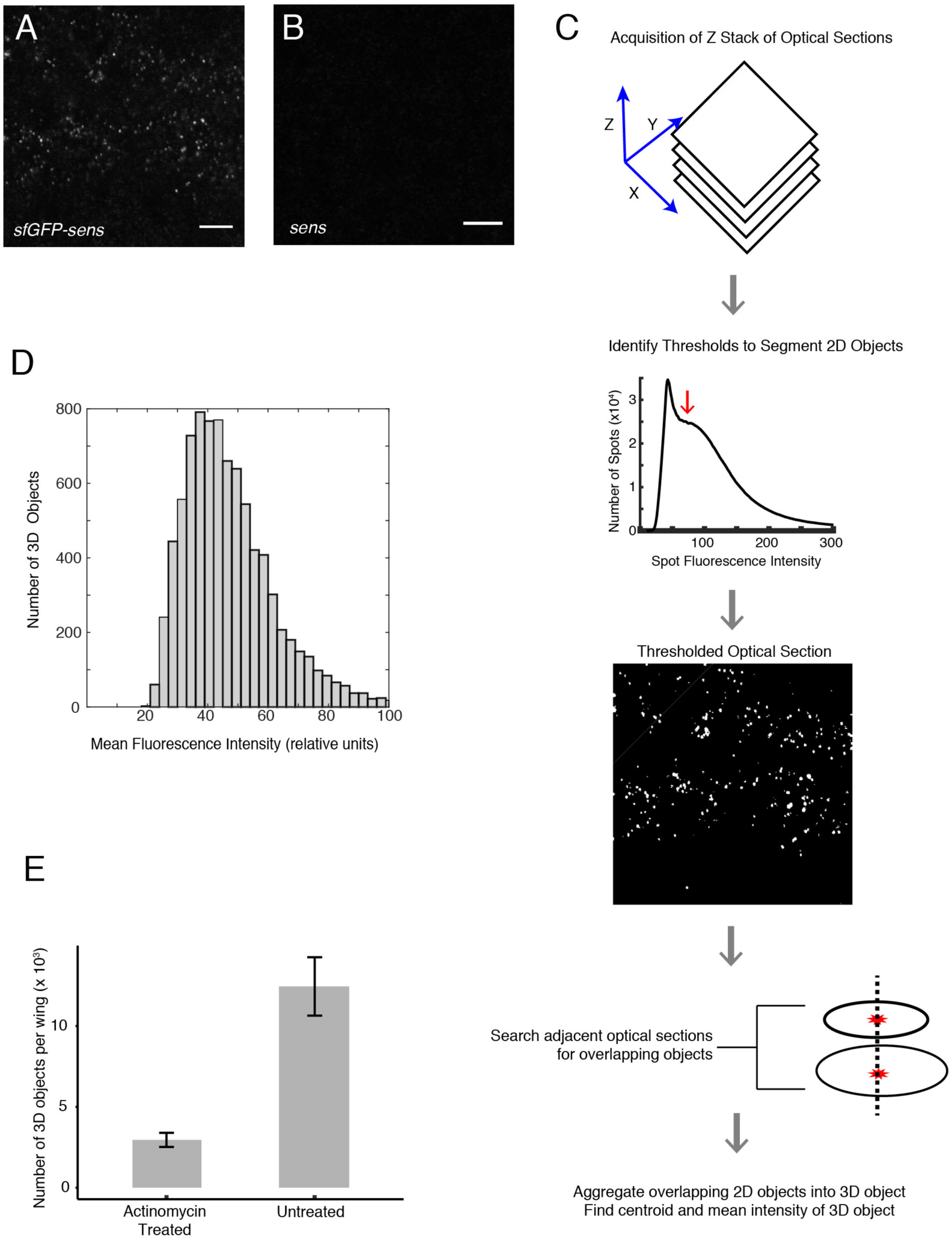
Development of smFISH imaging and analysis. **(A,B)** Representative optical sections of wing discs probed for sfGFP-containing mRNAs. Scale bars = 5 μm. **(A)** A disc from an animal with two copies of the *sfGFP-sens* transgene and two copies of the endogenous *sens^E1^* gene. **(B)** A disc from an animal with just two copies of the endogenous *sens^E1^* gene. **(C)** Imaging and analysis pipeline to quantify mRNAs as 3D fluorescent objects. **(D)** Distribution of mean fluorescence intensity for all identified 3D fluorescent objects from one wing disc expressing sfGFP-Sens mRNAs. **(E)** Average number of 3D fluorescent objects per imaged wing disc after a 30 minute treatment of the discs in actinomycin-D. Untreated discs were incubated in media for an identical period of time, and all discs were fixed and imaged for sfGFP-Sens mRNAs. Error bars are SEM.

**Figure 1 - figure supplement 2.**
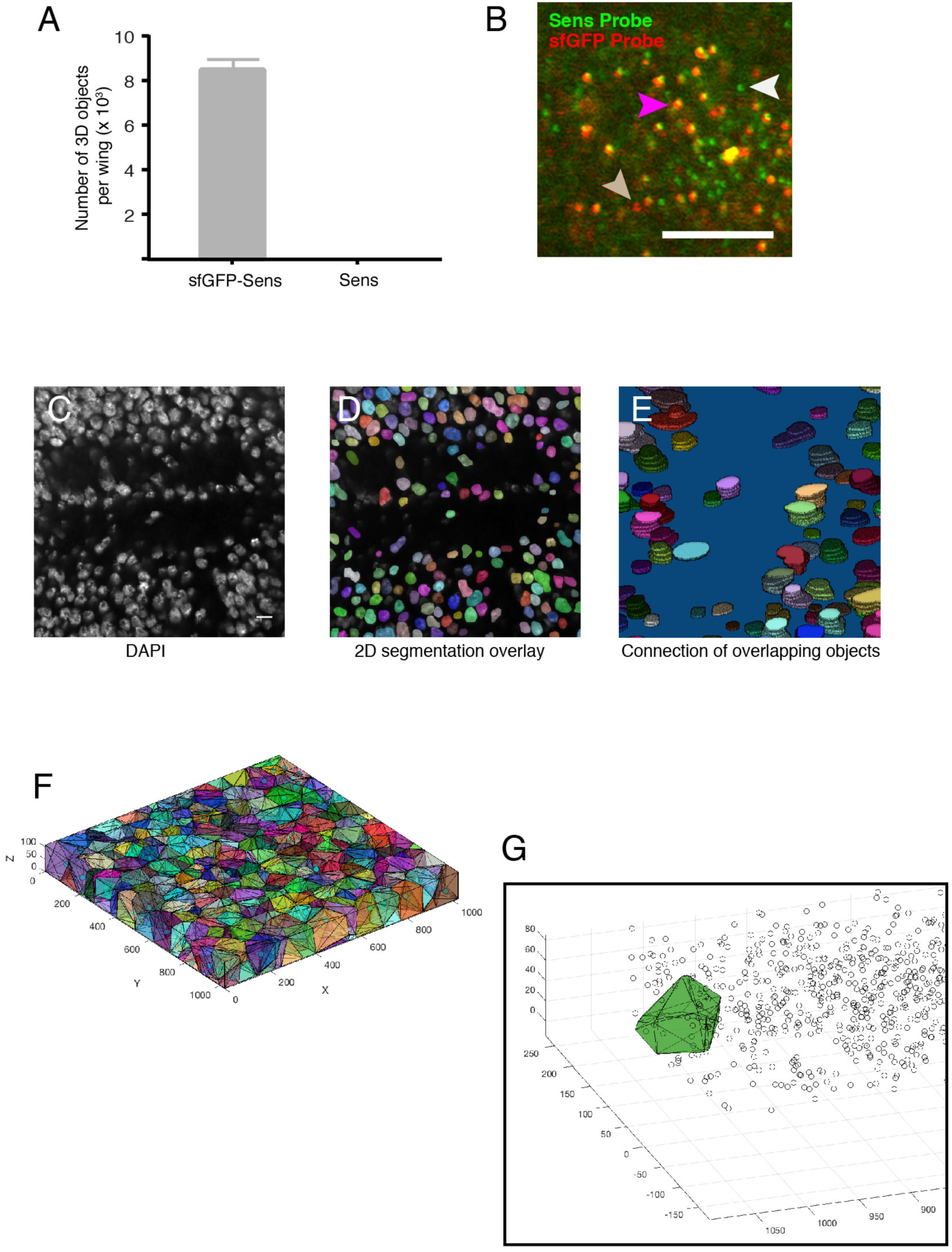
Determination of false-positive and false-negative rates for smFISH. **(A)** Wing discs were imaged and scored for 3D fluorescent objects using the sfGFP probe set. Discs were either from animals with two copies of the *sfGFP-sens* transgene and two copies of the endogenous *sens^E1^* gene, or from animals with just two copies of the endogenous *sens^E1^* gene. Error bars are SEM. **(B)** A representative optical section taken from a wing disc expressing the *sfGFP-sens* transgene and endogenous *sens^E1^* gene. The disc was probed for sfGFP (red) and Sens (green) RNA using independent probe sets. Spots that fluoresce both green and red are presumptive sfGFP-Sens mRNAs that have annealed to both probe sets (purple arrow). Spots that only fluoresce with the Sens probe set (white arrow) are presumptive Sens mRNAs that are generated from the endogenous *sens* gene. Although these *sens* alleles are mutant for protein output, they still produce mRNA. The occasional spot (beige arrow) that only fluoresces with the sfGFP probe set are presumptive sfGFP-Sens mRNAs that failed to hybridize with the Sens probe set. These are false-negatives. Scale bar = 5 μm. **(C-E)** Pipeline for 3D segmentation of cell nuclei. **(C)** An optical section showing DAPI fluorescence. **(D)** 2D segmentation of this image. **(E)** 3D segmentation by connecting 2D objects in neighboring sections that overlap with one another in the x-y plane. **(F)** 3D Voronoi tessellation of an image stack. The centroids of segmented nuclei (shown as circles) were used to tessellate the image stack, creating virtual cells. Cells are represented with different colors. Numbers in the x-y plane refer to pixel positions in the 1024 x 1024 sections. **(G)** An image stack showing the centroid positions of 3D mRNA objects as circles. One tessellated cell (green) is superimposed to show the mRNA objects that reside in space occupied by the tessellated cell. These mRNAs would be assigned to that particular cell. Shown is one stripe of sfGFP-Sens expressing cells on one side of the DV boundary marked by pixel position 0.

**Figure 1 - figure supplement 3.**
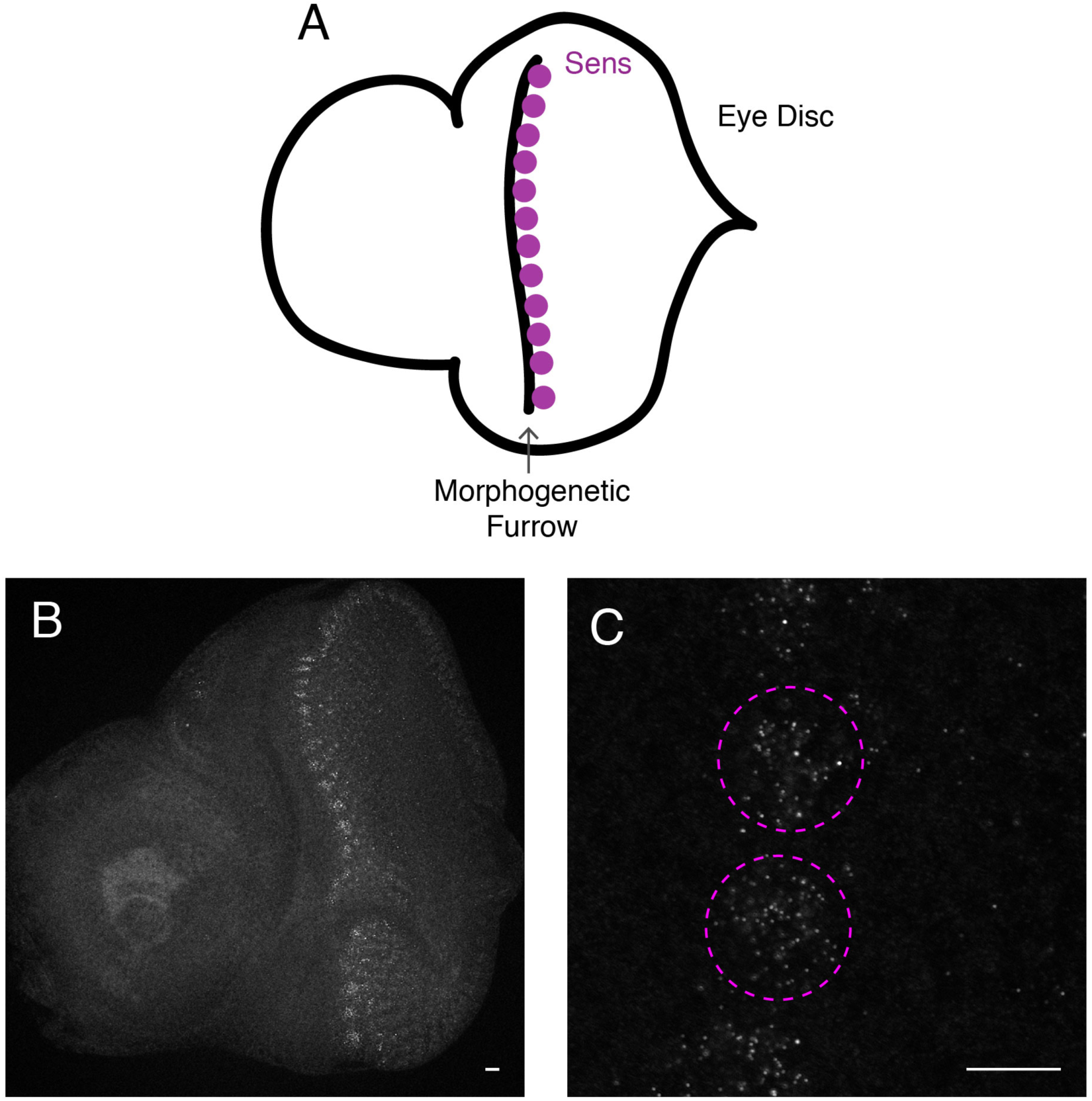
smFISH imaging of the eye imaginal disc. **(A)** Schematic of the eye antennal disc complex showing the approximate location of cells that express the *sens* gene. Anterior is to the left**. (B,C)** Optical sections through a representative eye antennal disc complex probed for sfGFP-Sens mRNAs by smFISH. Anterior is to the left. **(B)** Low magnification shows a vertical stripe of positive fluorescence that oscillates between clusters of high and low mRNA abundance. This is the pattern that has been reported for cells in the morphogenetic furrow (Nolo et al 2000). Scale bar = 5 μm. **(C)** Higher magnification of an optical section through the morphogenetic furrow showing two complete clusters of Sens-positive cells (dashed purple lines). Scale bar = 5 μm.

**Figure 3 - figure supplement 1.**
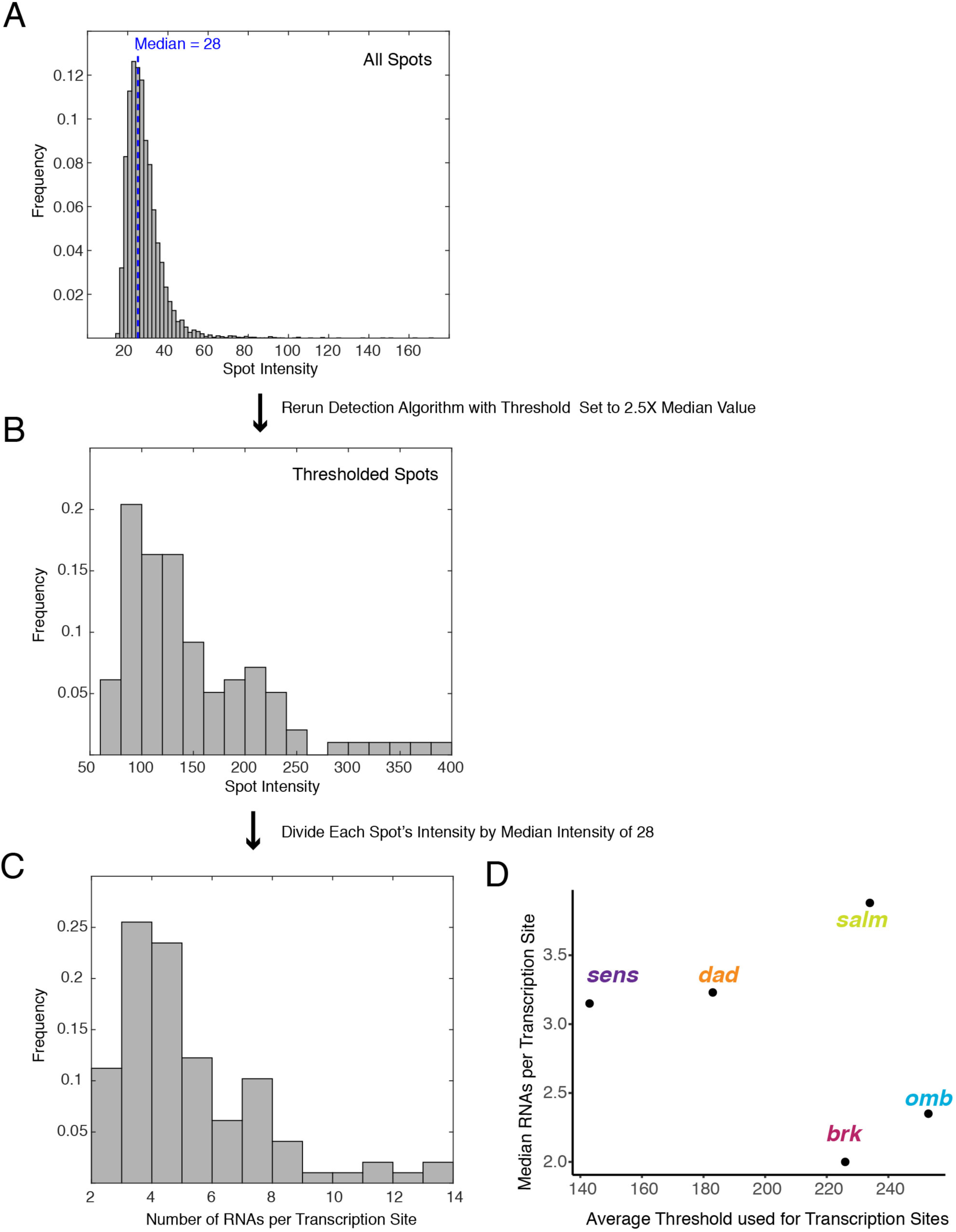
Detection of transcription sites and their quantification. **(A)** A representative frequency distribution of fluorescence intensity for 3D spots identified in one wing disc expressing *sfGFP-sens*. The median intensity is 28 units. **(B)** The same wing disc was reanalyzed for 3D spots but using a threshold of 70 units as a cutoff, below which spots are not counted. **(C)** The fluorescence intensity of each 3D spot in B is divided by the median intensity of 28 units to provide a normalized number of RNAs that are localized to that 3D spot. This is not an actual number of RNA molecules but the output from partially transcribed RNAs annealing to a variable number of probes depending on the composition of binding sites in the RNA composite. **(D)** The mean threshold used for transcription site identification for each data set plotted against the median normalized RNA molecules per transcription site for all transcription sites in that data set.

**Figure 3 - figure supplement 2.**
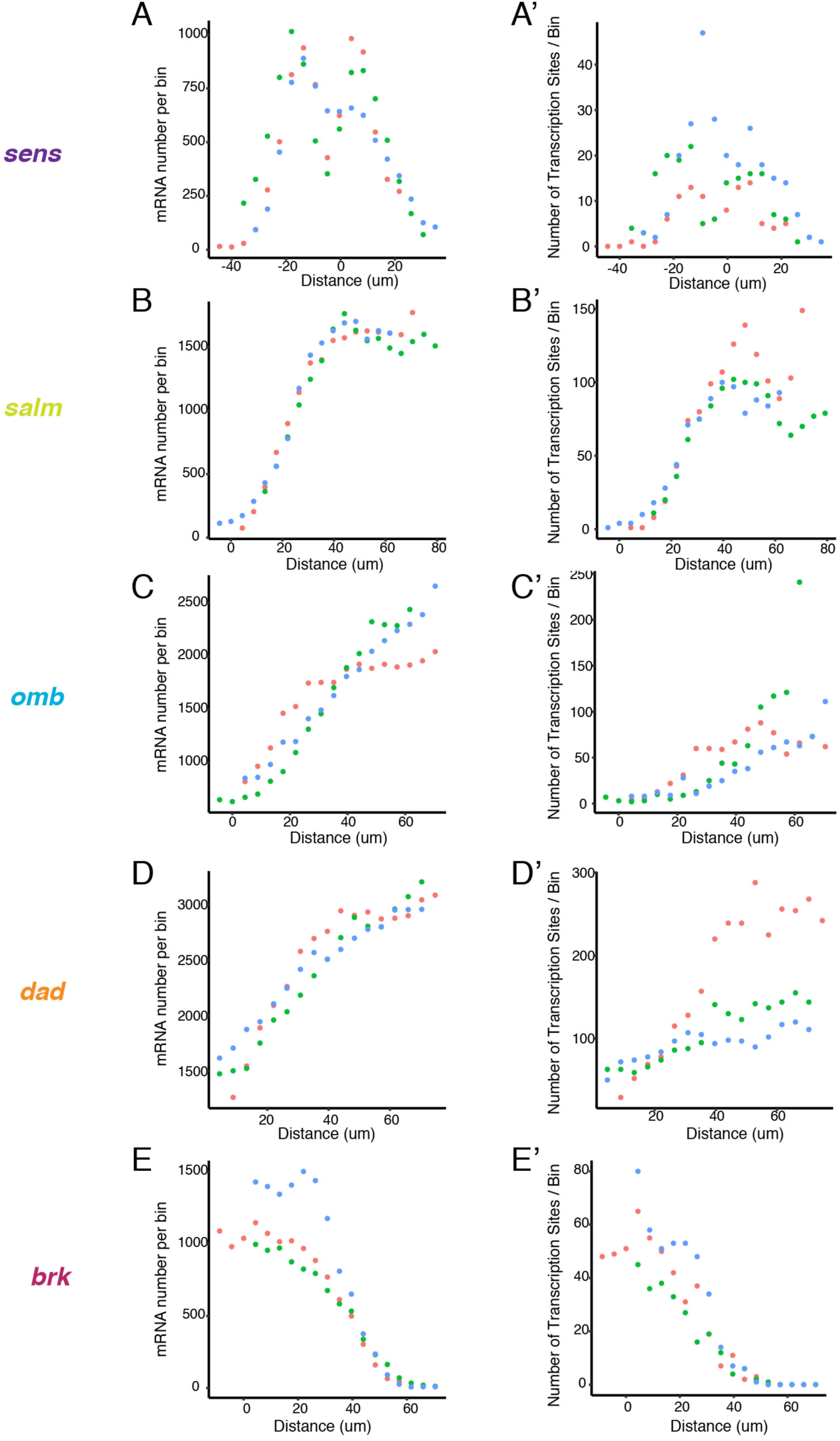
Transcription sites and mRNA patterns in unsegmented images. **(A,B,C,D,E)** Three discs were analyzed independently (green, blue, orange dots) for spots that corresponded to the mRNAs from *sens* **(A)**, *salm* **(B)**, *omb* **(C)**, *dad* **(D)** and *brk* **(E)**. Spots were binned according to their positions along the AP or DV axes, and total mRNAs per bin were plotted. Note the strong concordance of independent discs for all genes. **(A’,B’,C’,D’,E’)** The same three discs were analyzed independently (green, blue, orange dots) for spots that corresponded to transcription sites from *sens* **(A)**, *salm* **(B)**, *omb* **(C)**, *dad* **(D)** and *brk* **(E)**. Spots were binned according to their positions along the AP or DV axes, and total transcription sites per bin were plotted. Note the strong concordance of independent discs for all genes.

**Figure 6 - figure supplement 1.**
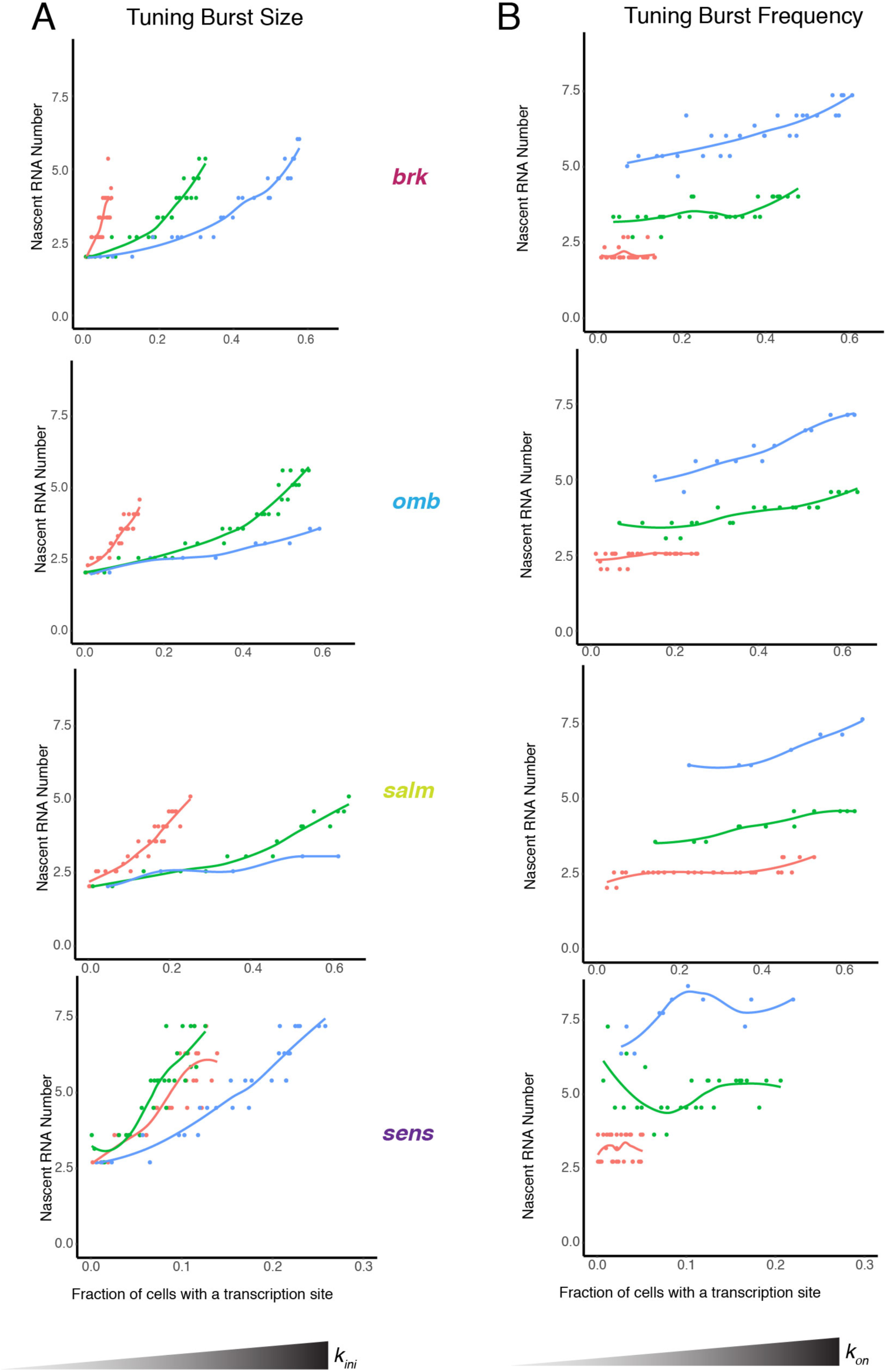
Modeling the relationship between average number of nascent RNAs in a transcription site and the probability of detecting a site for the *brk, omb, salm,* and *sens* genes. **(A)** Simulations are performed where the rate parameter *k_ini_* has been systematically varied so that burst size is variable. Resulting values for nascent RNA number and fraction of cells with a site are shown. Each datapoint is the average of 1,000 simulations. Simulations are repeated for three different values of *k_on_* to specifically set the burst frequency to 0.04, 0.2 and 0.4 min^-1^. **(B)** Simulations are performed where the rate parameter *k_on_* has been systematically varied so that burst frequency is variable. Resulting values for nascent RNA number and fraction of cells with a site are shown. Each datapoint is the average of 1,000 simulations. Simulations are repeated for three different values of *k_ini_* to specifically set the burst size to 1, 4 and 10.

